# A latent clinical-anatomical dimension relating metabolic syndrome to brain structure and cognition

**DOI:** 10.1101/2023.02.22.529531

**Authors:** Marvin Petersen, Felix Hoffstaedter, Felix L. Nägele, Carola Mayer, Maximilian Schell, D. Leander Rimmele, Birgit-Christiane Zyriax, Tanja Zeller, Simone Kühn, Jürgen Gallinat, Jens Fiehler, Raphael Twerenbold, Amir Omidvarnia, Kaustubh R. Patil, Simon B. Eickhoff, Götz Thomalla, Bastian Cheng

## Abstract

The link between metabolic syndrome (MetS) and neurodegenerative as well cerebrovascular conditions holds substantial implications for brain health in at-risk populations. This study elucidates the complex relationship between MetS and brain health by conducting a comprehensive examination of cardiometabolic risk factors, cortical morphology, and cognitive function in 40,087 individuals. Multivariate, data-driven statistics identified a latent dimension linking more severe MetS to widespread brain morphological abnormalities, accounting for up to 71% of shared variance in the data. This dimension was replicable across sub-samples. In a mediation analysis we could demonstrate that MetS-related brain morphological abnormalities mediated the link between MetS severity and cognitive performance in multiple domains. Employing imaging transcriptomics and connectomics, our results also suggest that MetS-related morphological abnormalities are linked to the regional cellular composition and macroscopic brain network organization. By leveraging extensive, multi-domain data combined with a dimensional stratification approach, our analysis provides profound insights into the association of MetS and brain health. These findings can inform effective therapeutic and risk mitigation strategies aimed at maintaining brain integrity.

## 1 Introduction

Metabolic syndrome (MetS) represents a cluster of cardiometabolic risk factors, including abdominal obesity, arterial hypertension, dyslipidemia, and insulin resistance [1]. With a prevalence of 23-35% in Western societies, it poses a considerable health challenge, promoting neurodegenerative and cerebrovascular diseases such as cognitive decline, dementia, and stroke [2–6]. As lifestyle and pharmacological interventions can modify the trajectory of MetS, advancing our understanding of its pathophysiological effects on brain structure and function as potential mediators of MetS-related neurological diseases is crucial to inform and motivate risk reduction strategies [7].

Magnetic resonance imaging (MRI) is a powerful non-invasive tool for examining the intricacies of neurological conditions in vivo. Among studies exploring MetS and brain structure, one of the most consistent findings has been alterations in cortical grey matter morphology [8]. Still, our understanding of the relationship between MetS and brain structure is constrained by several factors. To date, there have been only few studies on MetS effects on grey matter integrity that are well-powered [9–12]. The majority of analyses are based on small sample sizes and report effects only on global measures of brain morphology or a priori-defined regions of interest, limiting their scope [12–14]. As a result, reported effects are heterogeneous and most likely difficult to reproduce [15]. Existing large-scale analyses on the isolated effects of individual risk factors (such as hypertension or obesity) do not account for the high covariance of MetS components driven by interacting pathophysiological effects, which may prevent them from capturing the whole picture of MetS as a risk factor composite [16–19]. In addition, analyses addressing the complex interrelationship of MetS, brain structure and cognitive functioning by investigating them in conjunction are scarce [8]. Lastly, while previous studies adopted a case-control design treating MetS as a broad diagnostic category [10–12], a dimensional approach viewing MetS as a continuum could offer a more nuanced representation of the multivariate, continuous nature of the risk factor composite.

Despite reports on MetS effects on brain structure, the determinants and spatial effect patterns remain unclear. A growing body of evidence shows that spatial patterns of brain pathology are shaped by multi-scale neurobiological processes, ranging from the cellular level to regional dynamics to large-scale brain networks [20]. Accordingly, disease effects can not only be driven by local properties, when local patterns of tissue composition predispose individual regions to pathology, but also by topological properties of structural and functional brain networks [20,21]. Guided by these concepts, multi-modal and multi-scale analysis approaches could advance our understanding of the mechanisms influencing MetS effects on cortical morphology.

We argue that further research leveraging extensive clinical and brain imaging data is required to explore MetS effects on brain morphology. These examinations should integrate 1) a research methodology that strikes a balance between resolving the multivariate connection of MetS and brain structure while accounting for the high covariance of MetS components; 2) the recognition of impaired cognitive function as a pertinent consequence of MetS; and 3) the analysis of the spatial effect pattern of MetS and its possible determinants.

To meet these research needs, we investigated cortical thickness and subcortical volumetric measurements in a pooled sample of two large-scale population-based cohorts from the UK Biobank (UKB) and Hamburg City Health Study (HCHS) comprising in total 40,087 participants. Partial least squares correlation (PLS) analysis was employed to characterize MetS effects on regional brain morphology. PLS is especially suitable for this research task as it identifies overarching latent relationships by establishing a data-driven multivariate mapping between MetS components and brain morphometric indices. Furthermore, capitalizing on the cognitive phenotyping of both investigated cohorts, we examined the interrelation between MetS, cognitive function and brain structure in a mediation analysis. Finally, to uncover factors associated to brain region-specific MetS effects, we mapped local cellular as well as network topological attributes to observed MetS-associated cortical abnormalities. With this work, we aimed to advance the understanding of the fundamental principles underlying the neurobiology of MetS.

## 2 Materials and methods

### 2.1 Study population – the UK Biobank and Hamburg City Health Study

Here, we investigated cross-sectional clinical and imaging data from two large-scale population-based cohort studies: 1) the UK Biobank (UKB, n = 39,668, age 45-80 years; application number 41655) and 2) the Hamburg City Health Study (HCHS, n = 2637, age 45- 74 years) [22,23]. Both studies recruit large study samples with neuroimaging data alongside a detailed demographic and clinical assessment. Respectively, data for the first visit including a neuroimaging assessment were included. Individuals were excluded if they had a history or a current diagnosis of neurological or psychiatric disease. Field IDs of the used UKB variables are presented in *supplementary table S1*. UKB individuals were excluded based on the non-cancer illnesses codes (http://biobank.ndph.ox.ac.uk/showcase/coding.cgi?id=6). Excluded conditions were Alzheimer’s disease; alcohol, opioid and other dependencies; amyotrophic lateral sclerosis; brain injury; brain abscess; chronic neurological problem; encephalitis; epilepsy; haemorrhage; head injury; meningitis; multiple sclerosis; Parkinson’s disease; skull fracture. Same criteria were applied on HCHS individuals based on the neuroradiological evaluation and self-reported diagnoses variables. To enhance comparability to previous studies we supplemented a case-control analysis enabling to complement continuous multivariate statistical analyses by group statistics. Therefore, a MetS sample was identified based on the consensus definition of the International Diabetes Federation (*supplementary text S2*) and matched to a control cohort.

### 2.2 Ethics approval

The UKB was ethically approved by the North West Multi-Centre Research Ethics Committee (MREC). Details on the UKB Ethics and Governance framework are provided online (https://www.ukbiobank.ac.uk/media/0xsbmfmw/egf.pdf). The HCHS was approved by the local ethics committee of the Landesärztekammer Hamburg (State of Hamburg Chamber of Medical Practitioners, PV5131). Good Clinical Practice (GCP), Good Epidemiological Practice (GEP) and the Declaration of Helsinki were the ethical guidelines that governed the conduct of the HCHS [24]. Written informed consent was obtained from all participants investigated in this work.

### 2.3 Clinical assessment

In the UK Biobank, a battery of cognitive tests is administered, most of which represent shortened and computerized versions of established tests aiming for comprehensive and concise assessment of cognition [25]. From this battery we investigated tests for executive function and processing speed (Reaction Time Test, Symbol Digit Substitution Test, Tower Rearranging Test, Trail Making Tests A and B), memory (Numeric Memory Test, Paired Associate Learning Test, Prospective Memory Test) and reasoning (Fluid Intelligence Test, Matrix Pattern Completion Test). Detailed descriptions of the individual tests can be found elsewhere [26]. Furthermore, some tests (Matrix Pattern Completion Test, Numeric Memory Test, Paired Associate Learning Test, Symbol Digit Substitution Test, Trail Making Test and Tower Rearranging Test) are only administered to a subsample of the UKB imaging cohort explaining the missing test results for a subgroup of participants.

In the HCHS, cognitive testing was administered by a trained study nurse and included the Animal Naming Test, Trail Making Test A and B, Verbal Fluency and Word List Recall subtests of the Consortium to Establish a Registry for Alzheimer’s Disease Neuropsychological Assessment Battery (CERAD-Plus), as well as the Clock Drawing Test [27,28].

### 2.4 MRI acquisition

The full UKB neuroimaging protocol can be found online (https://biobank.ctsu.ox.ac.uk/crystal/crystal/docs/brain_mri.pdf) [22]. MR images were acquired on a 3-T Siemens Skyra MRI scanner (Siemens, Erlangen, Germany). T1-weighted MRI used a 3D MPRAGE sequence with 1-mm isotropic resolution with the following sequence parameters: repetition time = 2000 ms, echo time = 2.01 ms, 256 axial slices, slice thickness = 1 mm, and in-plane resolution = 1 x 1 mm. In the HCHS, MR images were acquired as well on a 3-T Siemens Skyra MRI scanner. Measurements were performed with a protocol as described in previous work [24]. In detail, for 3D T1-weighted anatomical images, rapid acquisition gradient-echo sequence (MPRAGE) was used with the following sequence parameters: repetition time = 2500 ms, echo time = 2.12 ms, 256 axial slices, slice thickness = 0.94 mm, and in-plane resolution = 0.83 × 0.83 mm.

### 2.5 Estimation brain morphological measures

To achieve comparability and reproducibility, the preconfigured and containerized CAT12 pipeline (CAT12.7 r1743; https://github.com/m-wierzba/cat-container) was employed for surface reconstruction and cortical thickness measurement building upon a projection-based thickness estimation method as well as computation of subcortical volumes [29]. Cortical thickness measures were normalized from individual to 32k fsLR surface space (conte69) to ensure vertex correspondence across subjects. Subcortical volumes were computed for the Melbourne Subcortex Atlas parcellation resolution 1 [30]. Volumetric measures for the anterior and posterior thalamus parcel were averaged to obtain a single measure for the thalamus. Individuals with a CAT12 image quality rating lower than 75% were excluded during quality assessment. To facilitate large-scale data management while ensuring provenance tracking and reproducibility, we employed the DataLad-based FAIRly big workflow for image data processing [31].

### 2.6 Statistical analysis

Statistical computations and plotting were performed in python 3.9.7 leveraging bctpy (v. 0.6.0), brainstat (v. 0.3.6), brainSMASH (v. 0.11.0), the ENIGMA toolbox (v. 1.1.3). matplotlib (v. 3.5.1), neuromaps (v. 0.0.1), numpy (v. 1.22.3), pandas (v. 1.4.2), pingouin (v. 0.5.1), pyls (v. 0.0.1), scikit-learn (v. 1.0.2), scipy (v. 1.7.3), seaborn (v. 0.11.2) as well as in matlab (v. 2021b) using ABAnnotate (v. 0.1.1).

#### 2.6.1 Partial least squares correlation analysis

To relate MetS components and cortical morphology, we performed a PLS using pyls (https://github.com/rmarkello/pyls). PLS identifies covariance profiles that relate two sets of variables in a data-driven double multivariate analysis [32]. Here, we related regional cortical thickness and subcortical volumes to clinical measurements of MetS components, i.e., obesity (waist circumference, hip circumference, waist-hip ratio, body mass index), arterial hypertension (systolic blood pressure, diastolic blood pressure), dyslipidemia (high density lipoprotein, low density lipoprotein, total cholesterol, triglycerides) and insulin resistance (HbA1c, non-fasting blood glucose). Before conducting the PLS, missing values were imputed via k-nearest neighbor imputation (n_neighbor_ = 4) with imputation only taking into account variables of the same group, i.e., MetS component variables were imputed based on the remaining MetS component data only and not based on demographic variables. To account for age, sex, education and cohort (UKB/HCHS) as potential confounds, they were regressed out of brain morphological and MetS component data.

We then performed PLS as described in previous work [33]. Methodological details are covered in *figure 1a* and *supplementary text S3*. Brain morphological measures were randomly permuted (n_random_ = 5000) to assess statistical significance of derived latent variables and their corresponding covariance profiles. Subject-specific PLS scores, including a clinical score and an imaging score, were computed. Higher scores indicate stronger adherence to the respective covariance profiles: a high clinical score signifies pronounced expression of the clinical profile, and a high imaging score reflects marked adherence to the brain morphological profile. Bootstrap resampling (n_bootstrap_ = 5000) was performed to assess the contribution of individual variables to the imaging-clinical relationship. Confidence intervals (95%) of singular vector weights were computed for clinical variables to assess the significance of their contribution. To estimate contributions of brain regions, bootstrap ratios were computed as the singular vector weight divided by the bootstrap-estimated standard error. A high bootstrap ratio is indicative of a region’s contribution, as a relevant region shows a high singular vector weight alongside a small standard error implying stability across bootstraps. The bootstrap ratio equals a z-score in case of a normally distributed bootstrap. Hence, brain region contributions were considered significant if the bootstrap ratio was >1.96 or <-1.96 (95% confidence interval). Overall model robustness was assessed via a 10-fold cross-validation by correlating out-of-sample PLS scores within each fold.

**Figure 1.**
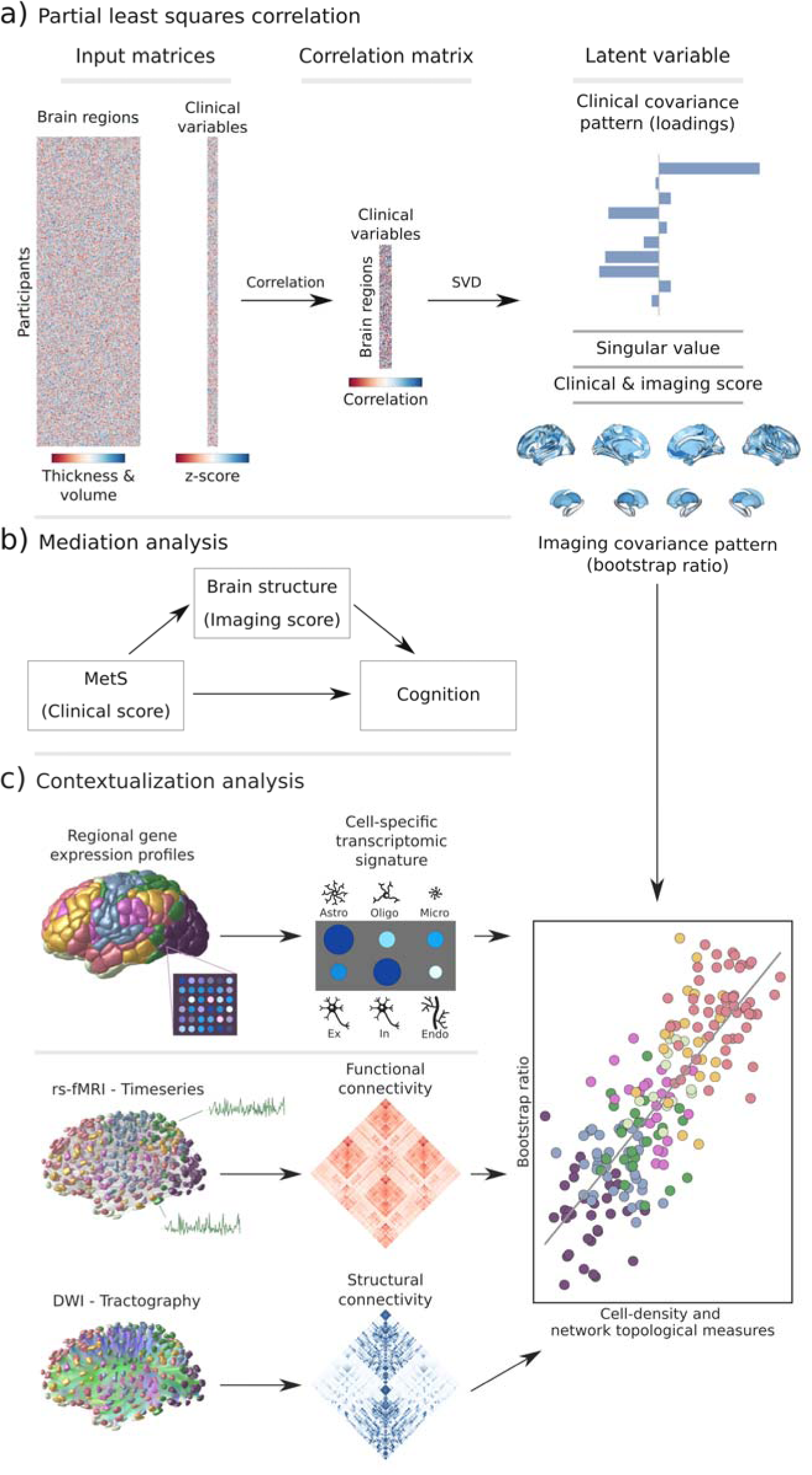
Methodology. a) Illustration of the partial least squares correlation analysis. Starting from two input matrices containing per-subject information of regional morphological measures as well as clinical data (demographic and MetS-related risk factors) a correlation matrix is computed. This matrix is subsequently subjected to singular value decomposition resulting in a set of mutually orthogonal latent variables. Latent variables each consist of a left singular vector (here, clinical covariance profile), singular value and right singular vector (here, imaging covariance profile). In addition, subject-specific clinical and imaging scores are computed. b) The interplay between MetS, brain structure and cognition was investigated in a post-hoc mediation analysis. We tested whether the relationship between the clinical score, representing MetS severity, and different cognitive test performances was statistically mediated by the imaging score. c) Contextualization analysis. Upper row: based on microarray gene expression data, the densities of different cell populations across the cortex were quantified. Middle and lower row: based on functional and structural group-consensus connectomes based on data from the Human Connectome Project, metrics of functional and structural brain network topology were derived. Cell density as well as connectomic measures were related to the bootstrap ratio via spatial correlations. Modified from Petersen et al. and Zeighami et al. [33,82]. Abbreviations: Astro – astrocytes; DWI – diffusion-weighted magnetic resonance imaging; Endo – endothelial cells; Ex – excitatory neuron populations (Ex1-8); In – inhibitory neuron populations (In1-8); Micro – microglia; Oligo – oligodendrocytes; rs-fMRI – resting-state functional magnetic resonance imaging; SVD – singular value decomposition.

#### 2.6.2 Mediation analysis

In a post-hoc mediation analysis, we investigated how the subject-specific clinical PLS score of the first latent variable, reflecting the degree of an individual’s expression of the identified MetS risk profile, relates to cognitive test outcomes, and whether this relationship is influenced by the imaging PLS score of the first latent variable, which represents the degree of brain morphological differences (*figure 1b*). This analysis allows to separate the total effect of the clinical PLS score on cognitive performance into: (1) a direct effect (the immediate impact of clinical scores on cognition), and (2) an indirect effect (the portion influenced by the imaging PLS score). This approach helps to disentangle the complex interplay between MetS and cognitive function by examining the role of brain structural effects as a potential intermediary. We considered an indirect effect as mediating if there was a significant association between the clinical and imaging PLS scores, the imaging PLS score was significantly associated to the cognitive outcome, and if the link between clinical scores and cognitive outcomes weakened (partial mediation) or became insignificant (full mediation) after accounting for imaging scores. The significance of mediation was assessed using bootstrapping (n_bootstrap_=5000), with models adjusted for age, sex, and education. To obtain standardized estimates, mediation analysis inputs were z-scored beforehand. Given the variation in cognitive test batteries between the UKB and HCHS cohorts, only individuals with results from the respective tests were considered in each mediation analysis. To account for the different versions of the Trail Making Tests A and B used in both cohorts, test results were harmonized through z-scoring within the individual subsamples before pooled z-scoring.

#### 2.6.3 Contextualization analysis

We investigated the link of MetS and regional brain morphological measurements in the context of cell-specific gene expression profiles and structural and functional brain network characteristics (*figure 1c*). Therefore, we used the Schaefer-parcellated (400×7 and 100×7, v.1) bootstrap ratio map and related it to indices representing different gene expression and network topological properties of the human cortex via spatial correlations (Spearman correlation, r_sp_) on a group-level [34].

Virtual histology analysis. We performed a virtual histology analysis leveraging gene transcription information to quantify the density of different cell populations across the cortex employing the ABAnnotate toolbox [35,36]. Genes corresponding with specific cell populations of the central nervous system were identified based on a classification derived from single nucleus-RNA sequencing data [37]. The gene-celltype mapping is provided by the PsychENCODE database (http://resource.psychencode.org/Datasets/Derived/SC_Decomp/DER-19_Single_cell_markergenes_TPM.xlsx) [38]. The abagen toolbox (v. 0.1.3) was used to obtain regional microarray expression data of these genes for Schaefer100×7 parcels based on the Allen Human Brain Atlas (AHBA) [39]. The Schaefer100×7 atlas was used as it better matches the sampling density of the AHBA eventually resulting in no parcels with missing values. Regional expression patterns of genes corresponding to astrocytes, endothelial cells, excitatory neuron populations (Ex1-8), inhibitory neuron populations (In1-8), microglia, and oligodendrocytes were extracted. Instead of assessing the correspondence between MetS effects and the expression pattern of each gene directly, we employed ensemble-based gene category enrichment analysis (GCEA) [40]. This approach represents a modification to customary GCEA addressing the issues of gene-gene dependency through within-category co-expression which is caused by shared spatial embedding as well as spatial autocorrelation of cortical transcriptomics data. In brief, gene transcription indices were averaged within categories (here cell populations) and spatially correlated with the bootstrap ratio map. Statistical significance was assessed by comparing the empirical correlation coefficients against a null distribution derived from surrogate maps with preserved spatial embedding and autocorrelation computed via a spatial lag model [41]. Further details on the processing steps covered by ABAnnotate can be found elsewhere (https://osf.io/gcxun) [42].

Brain network topology. To investigate the cortical MetS effects pattern in the context of brain network topology, three connectivity metrics were leveraged based on data from structural and functional brain imaging: weighted degree centrality, neighborhood abnormality as well as macroscale functional connectivity gradients as described previously [33]. These were computed based on functional and structural consensus connectomes on group-level derived from the Human Connectome Project Young Adults dataset comprised in the ENIGMA toolbox [43,44]. Computation and derivation of the metrics are described in the *supplementary text S4*. For this analysis, statistical significance of spatial correlations was assessed via spin permutations (n = 1,000) which represent a null model preserving the inherent spatial autocorrelation of cortical information [45]. Spin permutations are performed by projecting parcel-wise data onto a sphere which then is randomly rotated. After rotation, information is projected back on the surface and a permuted r_sp_ is computed. A p-value is computed comparing the empirical correlation coefficient to the permuted distribution. To assure that our results do not depend on null model choice, we additionally tested our results against a variogram-based null model implemented in the brainSMASH toolbox (https://github.com/murraylab/brainsmash) as well as a network rewiring null model with preserved density and degree sequence [46,47].

All p-values resulting from both contextualization analyses were FDR-corrected for multiple comparisons. As we conducted this study mindful of the reuse of our resources, the MetS effect maps are provided as separate supplementary files to enable further analyses.

#### 2.6.4 Sensitivity analyses

For a sensitivity analysis, we reperformed the PLS separately within the UKB and HCHS cohorts. In contrast to the PLS main analysis, in these subset specific PLS analyses cognitive test performances were also incorporated as clinical variables as cognitive batteries were subset specific. This approach was employed to evaluate the stability of the results and to determine if cognitive tests contribute to the latent variables.

To test whether the PLS indeed captures the link of MetS and brain morphology, we conducted a group comparison as in previous studies of MetS. Besides descriptive group statistics, the cortical thickness of individuals with MetS and matched controls was compared on a surface vertex-level leveraging the BrainStat toolbox (v 0.3.6, https://brainstat.readthedocs.io/) [48]. A general linear model was applied correcting for age, sex, education and cohort effects. Vertex-wise p-values were FDR-corrected for multiple comparisons. To demonstrate the correspondence between the t-statistic and cortical bootstrap ratio maps, we related them via spatial correlation analyses. The t-statistic map was also used for sensitivity analysis of the virtual histology analysis and brain network contextualization.

To ensure that the brain network contextualization results were not biased by the connectome choice, we reperformed the analysis with structural and functional group consensus connectomes based on resting-state functional and diffusion-weighted MRI data from the HCHS. The corresponding connectome reconstruction approaches were described elsewhere [33].

## 3 Results

### 3.1 Sample characteristics

Application of exclusion criteria and quality assessment ruled out 2,188 UKB subjects and 30 HCHS subjects resulting in a final analysis sample of 40,087 individuals. For a flowchart providing details on the sample selection procedure please refer to *supplementary figure S5*. Descriptive statistics are listed in *table 1*. To sensitivity analyze our results, as well as to facilitate the comparison with previous reports which primarily rely on a case-control design, we supplemented group statistics comparing individuals with clinically defined MetS and matched controls, where applicable. Corresponding group analysis results are described in more detail in *supplementary materials S6-12*.

**Table 1.**
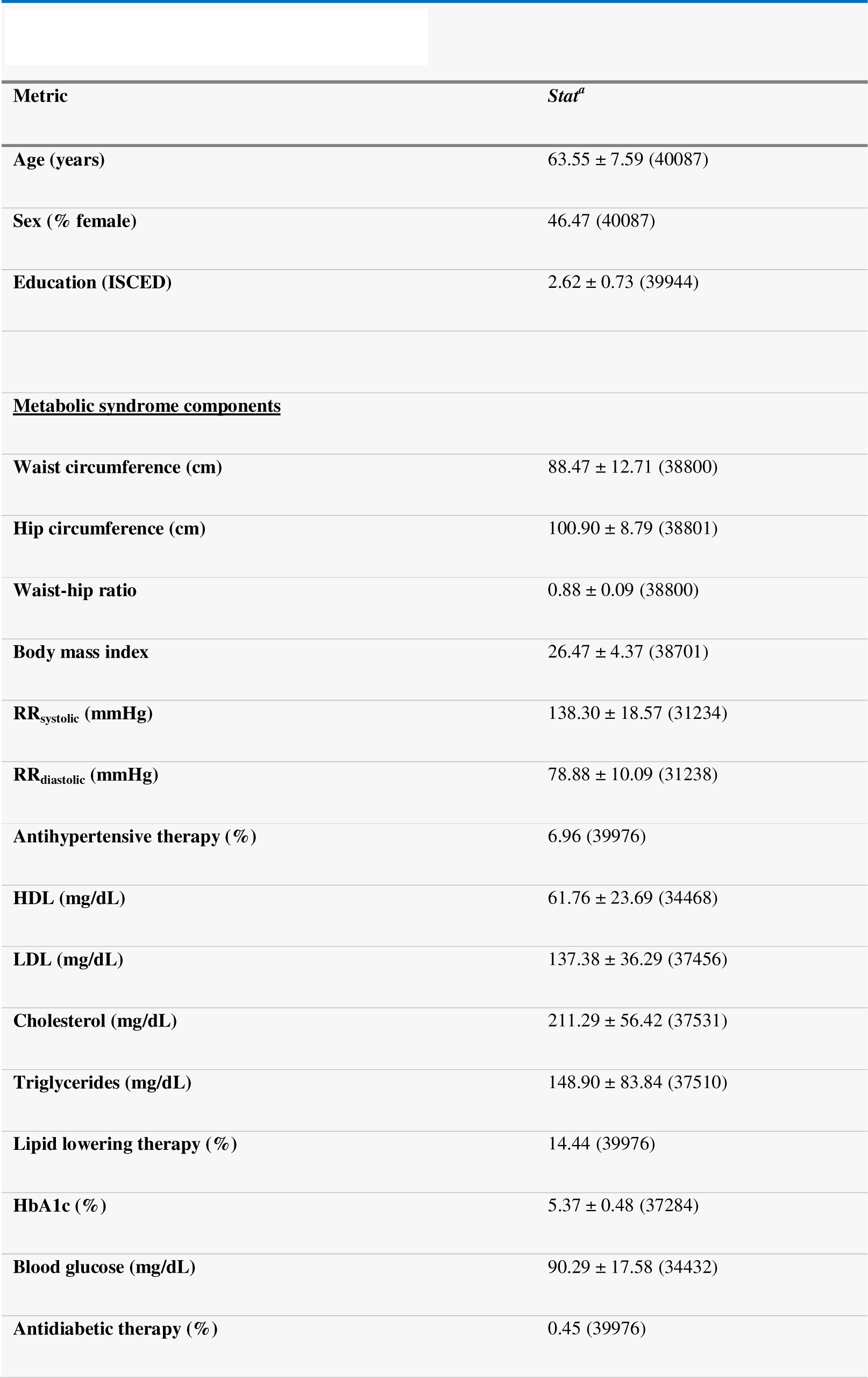
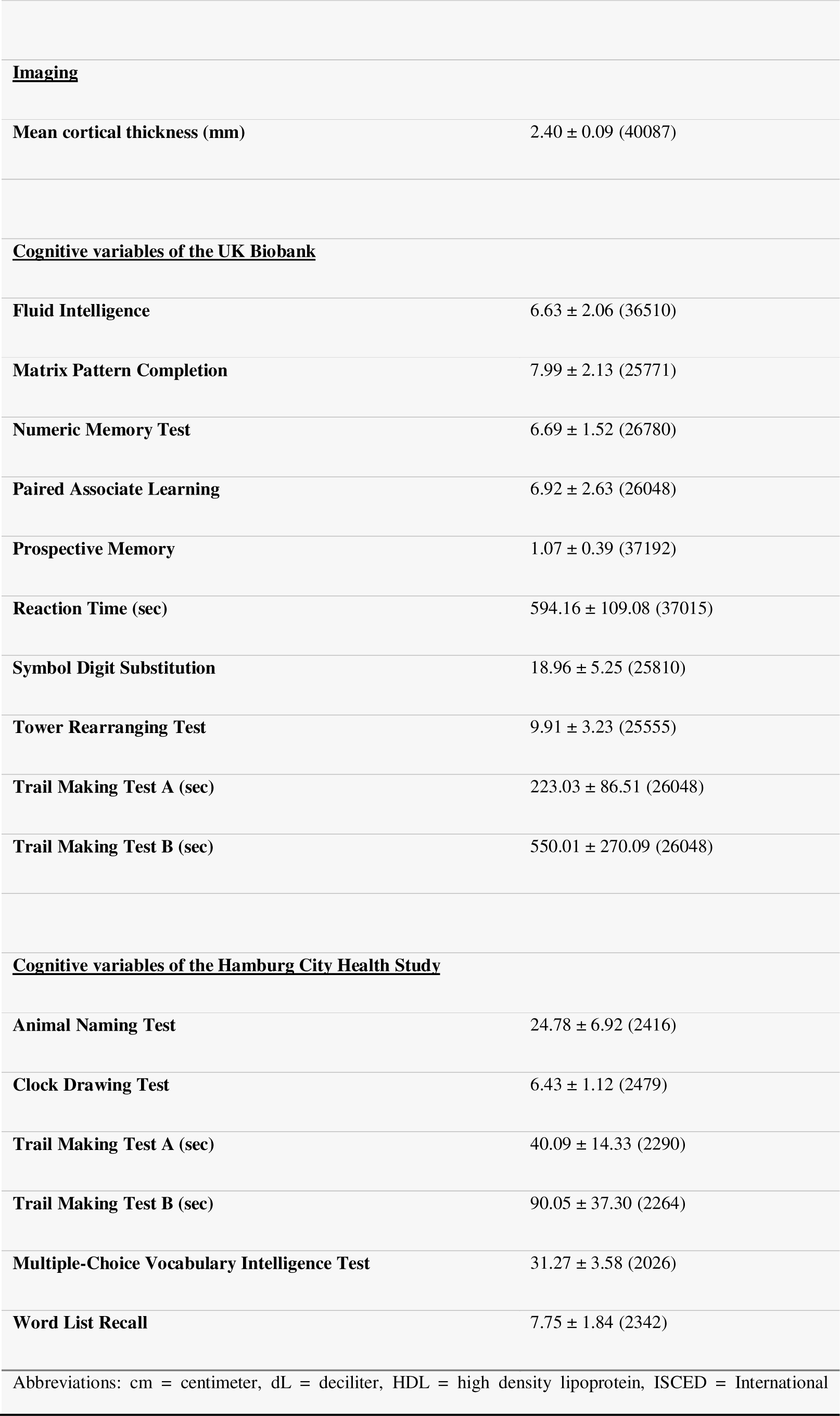
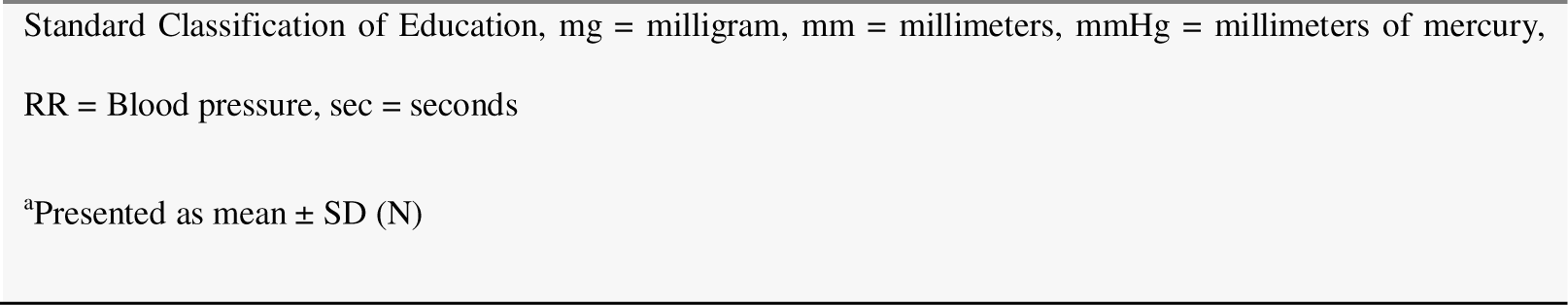
Descriptive statistics UKB and HCHS.

| Table 1. Descriptive statistics UKB and HCHS |  |
| --- | --- |
| Metric | Stat<sup>a</sup> |
| Age (years) | 63.55 ± 7.59 (40087) |
| Sex (% female) | 46.47 (40087) |
| Education (ISCED) | 2.62 ± 0.73 (39944) |
| **<u>Metabolic syndrome components</u>** |  |
| Waist circumference (cm) | 88.47 ± 12.71 (38800) |
| Hip circumference (cm) | 100.90 ± 8.79 (38801) |
| Waist-hip ratio | 0.88 ± 0.09 (38800) |
| Body mass index | 26.47 ± 4.37 (38701) |
| RR<sub>systolic</sub> (mmHg) | 138.30 ± 18.57 (31234) |
| RR<sub>diastolic</sub> (mmHg) | 78.88 ± 10.09 (31238) |
| Antihypertensive therapy (%) | 6.96 (39976) |
| HDL (mg/dL) | 61.76 ± 23.69 (34468) |
| LDL (mg/dL) | 137.38 ± 36.29 (37456) |
| Cholesterol (mg/dL) | 211.29 ± 56.42 (37531) |
| Triglycerides (mg/dL) | 148.90 ± 83.84 (37510) |
| Lipid lowering therapy (%) | 14.44 (39976) |
| HbA1c (%) | 5.37 ± 0.48 (37284) |
| Blood glucose (mg/dL) | 90.29 ± 17.58 (34432) |
| Antidiabetic therapy (%) | 0.45 (39976) |
| **<u>Imaging</u>** |  |
| Mean cortical thickness (mm) | 2.40 ± 0.09 (40087) |
| **<u>Cognitive variables of the UK Biobank</u>** |  |
| Fluid Intelligence | 6.63 ± 2.06 (36510) |
| Matrix Pattern Completion | 7.99 ± 2.13 (25771) |
| Numeric Memory Test | 6.69 ± 1.52 (26780) |
| Paired Associate Learning | 6.92 ± 2.63 (26048) |
| Prospective Memory | 1.07 ± 0.39 (37192) |
| Reaction Time (sec) | 594.16 ± 109.08 (37015) |
| Symbol Digit Substitution | 18.96 ± 5.25 (25810) |
| Tower Rearranging Test | 9.91 ± 3.23 (25555) |
| Trail Making Test A (sec) | 223.03 ± 86.51 (26048) |
| Trail Making Test B (sec) | 550.01 ± 270.09 (26048) |
| **<u>Cognitive variables of the Hamburg City Health Study</u>** |  |
| Animal Naming Test | 24.78 ± 6.92 (2416) |
| Clock Drawing Test | 6.43 ± 1.12 (2479) |
| Trail Making Test A (sec) | 40.09 ± 14.33 (2290) |
| Trail Making Test B (sec) | 90.05 ± 37.30 (2264) |
| Multiple-Choice Vocabulary Intelligence Test | 31.27 ± 3.58 (2026) |
| Word List Recall | 7.75 ± 1.84 (2342) |
| Abbreviations: cm = centimeter, dL = deciliter, HDL = high density lipoprotein, ISCED = International |  |

Standard Classification of Education, mg = milligram, mm = millimeters, mmHg = millimeters of mercury,
RR = Blood pressure, sec = seconds
<sup>a</sup>Presented as mean ± SD (N)

### 3.2 Partial least squares correlation analysis

We investigated the relationship between brain morphological and clinical measures of MetS (abdominal obesity, arterial hypertension, dyslipidemia, insulin resistance) in a PLS considering all individuals from both studies (n=40,087). By this, we aimed to detect the continuous effect of any MetS component independent from a formal binary classification of MetS (present / not present). A correlation matrix relating all considered MetS component measures is displayed in *supplementary figure S13*. Before conducting the PLS, brain morphological and clinical data were deconfounded for age, sex, education and cohort effects.

PLS identified seven significant latent variables which represent clinical-anatomical dimensions relating MetS components to brain morphology (*supplementary table S14*). The first latent variable explained 71.20% of shared variance and was thus further investigated (*figure 2a*). Specifically, the first latent variable corresponded with a covariance profile of lower severity of MetS (*figure 2c*; loadings [95% confidence interval]; waist circumference: -.230 [-.239,-.221], hip circumference: -.187 [-.195,-.178], waist-hip ratio: -.167 [-.176,-.158], body mass index: -.234 [-.243,-.226], systolic blood pressure: -.089 [-.098,-.080], diastolic blood pressure: -.116 [-.125,-.107], high density lipoprotein: .099 [.090,.108], low density lipoprotein: -.013 [-.022,-.004], total cholesterol: .003 [-.006,.012], triglycerides: -.102 [-.111,-.092], HbA1c: -.064 [-.073,-0.54], glucose: -.049 [-.058,-.039]). Notably, the obesity-related measures showed the strongest contribution to the covariance profile as indicated by the highest loading to the latent variable. Age (<.001 [-.009,.009]), sex (<.001 [-.009,.009]), education (<.001 [-.009,.009]) and cohort (<-.001 [-.008,.007]) did not significantly contribute to the latent variable, which is compatible with sufficient effects of deconfounding. Details on the second latent variable which explained 22.33% of shared variance are provided in *supplementary figure S1*5. In brief, it predominantly related lower HbA1c and blood glucose to higher thickness and volume in lateral frontal, posterior temporal, parietal and occipital regions and vice versa.

**Figure 2.**
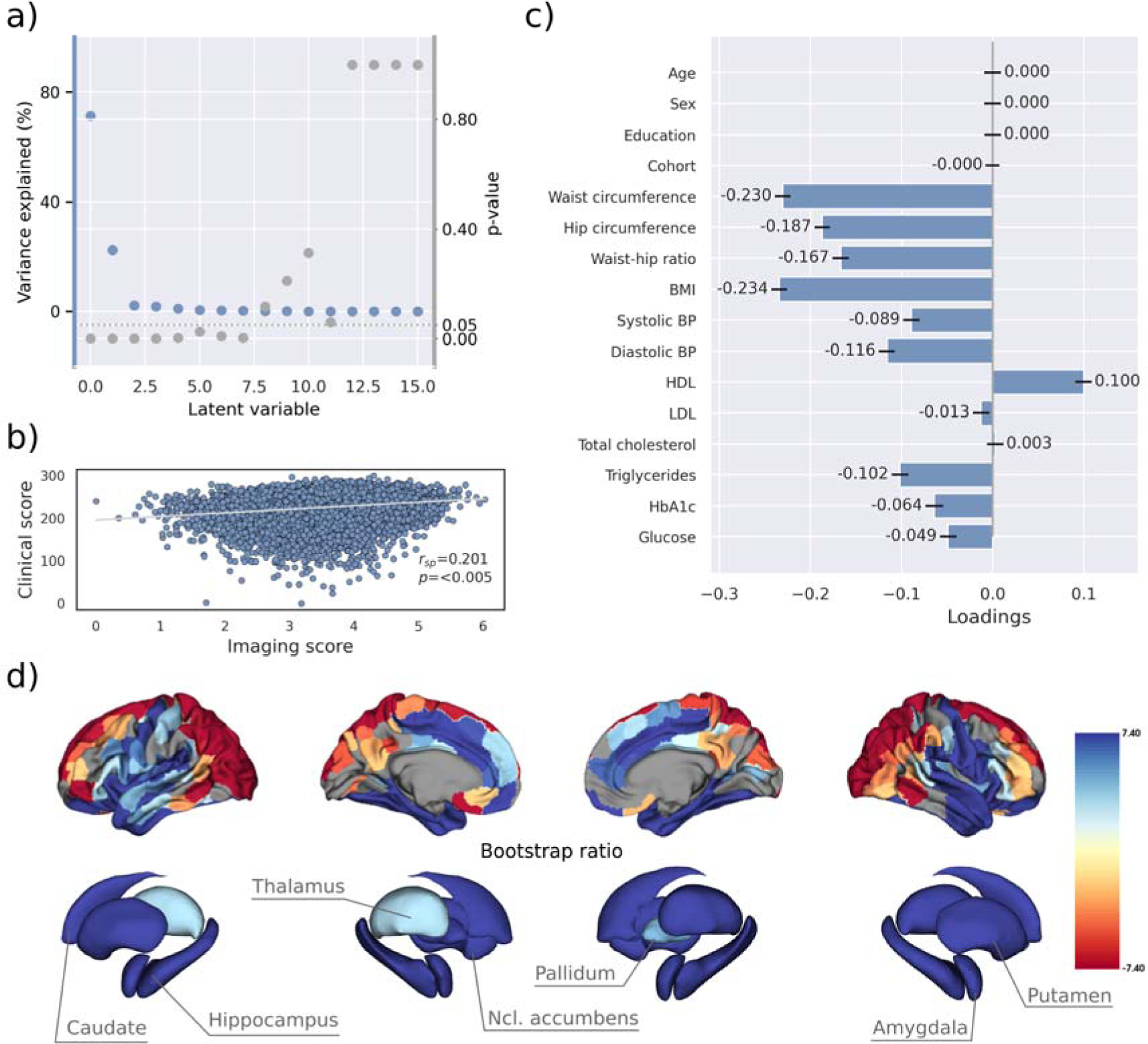
Partial least squares (PLS) analysis. a) Explained variance and p-values of latent variables. b) Scatter plot relating subject-specific clinical and imaging PLS scores. Higher scores indicate higher adherence to the respective covariance profile. c) Clinical covariance profile. 95% confidence intervals were calculated via bootstrap resampling. Note that confound removal for age, sex, education and cohort was performed prior to the PLS. d) Imaging covariance profile represented by bootstrap ratio. A high positive or negative bootstrap ratio indicates high contribution of a brain region to the overall covariance profile. Vertices with a significant bootstrap ratio (> 1.96 or < ™1.96) are highlighted by colors. Abbreviations: - Spearman correlation coefficient.

Bootstrap ratios 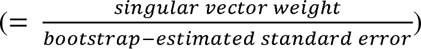 were computed to identify brain regions with a significant contribution to the covariance profile (see Methods). Cortical thickness in orbitofrontal, lateral prefrontal, insular, anterior cingulate and temporal areas as well as all volumes of all investigated subcortical regions contributed positively to the covariance profile as indicated by a positive bootstrap ratio (*figure 2d*). Thus, a higher cortical thickness and subcortical volume in these areas corresponded with less obesity, hypertension, dyslipidemia and insulin resistance and vice versa, i.e., lower cortical thickness and subcortical volumes with increased severity of MetS. A negative bootstrap ratio was found in superior frontal, parietal and occipital regions indicating that a higher cortical thickness in these regions corresponded with a more pronounced expression of MetS components. This overall pattern was confirmed via conventional, vertex-wise group comparisons of cortical thickness measurements based on the binary classification of individuals with MetS and matched controls (*supplementary figure S12*) as well as subsample analyses considering the UKB and HCHS participants independently (*supplementary figure S16-17*). The correlation matrix of all spatial effect maps investigated in this study (bootstrap ratio and Schaefer400-parcellated t-statistic from group comparisons) is visualized in *supplementary figure S18*. All derived effect size maps were significantly correlated (r_sp_ =.67 - .99, p_FDR_ < .05) [34].

Subject-specific imaging and clinical scores for the first latent variable were computed. These scores indicate to which degree an individual expresses the corresponding covariance profiles. By definition, the scores are correlated (r_sp_ = .201, p<.005, *figure 2b*) indicating that individuals exhibiting the clinical covariance profile (severity of MetS components) also express the brain morphological pattern. This relationship was robust across a 10-fold cross-validation (avg. r_sp_ = .19, *supplementary table S19*).

These results were consistent in separate PLS analyses for both the UKB and HCHS samples, as displayed in *supplementary figures S16 and S17*. In these subset-specific analyses, cognitive test performances significantly contributed to the first latent variable when included in the PLS. Consequently, the first latent variable associated more severe MetS with both brain morphological abnormalities and poorer cognitive performance.

### 3.3 Mediation analysis of cognitive outcomes

To gain a better understanding of the link between MetS, brain morphology, and cognitive function, we performed a mediation analysis on cognitive test results and subject-specific PLS scores. Therefore, we investigated whether the imaging PLS score (representing MetS-related brain structural abnormalities) acts as a mediator in the relationship between the clinical PLS score (representing MetS severity) and cognitive test performances. Importantly, scores of the main PLS analysis, which did not include cognitive measures, were considered. The corresponding path plots are shown in *figure 3*. The imaging score was found to fully mediate the relationship of the clinical score and results of the Trail Making Test B (ab = -.011, *P_FDR_* < .001; c’ = -.012, *P_FDR_* = .072; c = ™0.023, *P_FDR_* < 0.001), Fluid Intelligence Test (ab = .017, *P_FDR_* < .001; c’ = .011, *P_FDR_* = .072; c = .028, *P_FDR_* < .001) as well as Matrix Pattern Completion Test (ab = .015, *P_FDR_* < .001; c’ = .010, *P_FDR_* = .172; c = .025, *P_FDR_* <.001). Further, the imaging score partially mediated the relationship of the clinical score and results of the Symbol Digit Substitution Test (ab = .010, *P_FDR_* < .001; c’ = .036, *P_FDR_* < .001; c = 0.046, *P_FDR_* < 0.001), Numeric Memory Test (ab = .014, *P_FDR_* < .001; c’ = .044, *P_FDR_* <.001; c = 0.058, *P_FDR_* < 0.001) and Paired Associate Learning Test (ab = .015, *P_FDR_* < .001; c’ = .044, *P_FDR_* < .001; c = 0.059, *P_FDR_* < 0.001). For the remaining cognitive tests, no significant mediation was found.

**Figure 3.**
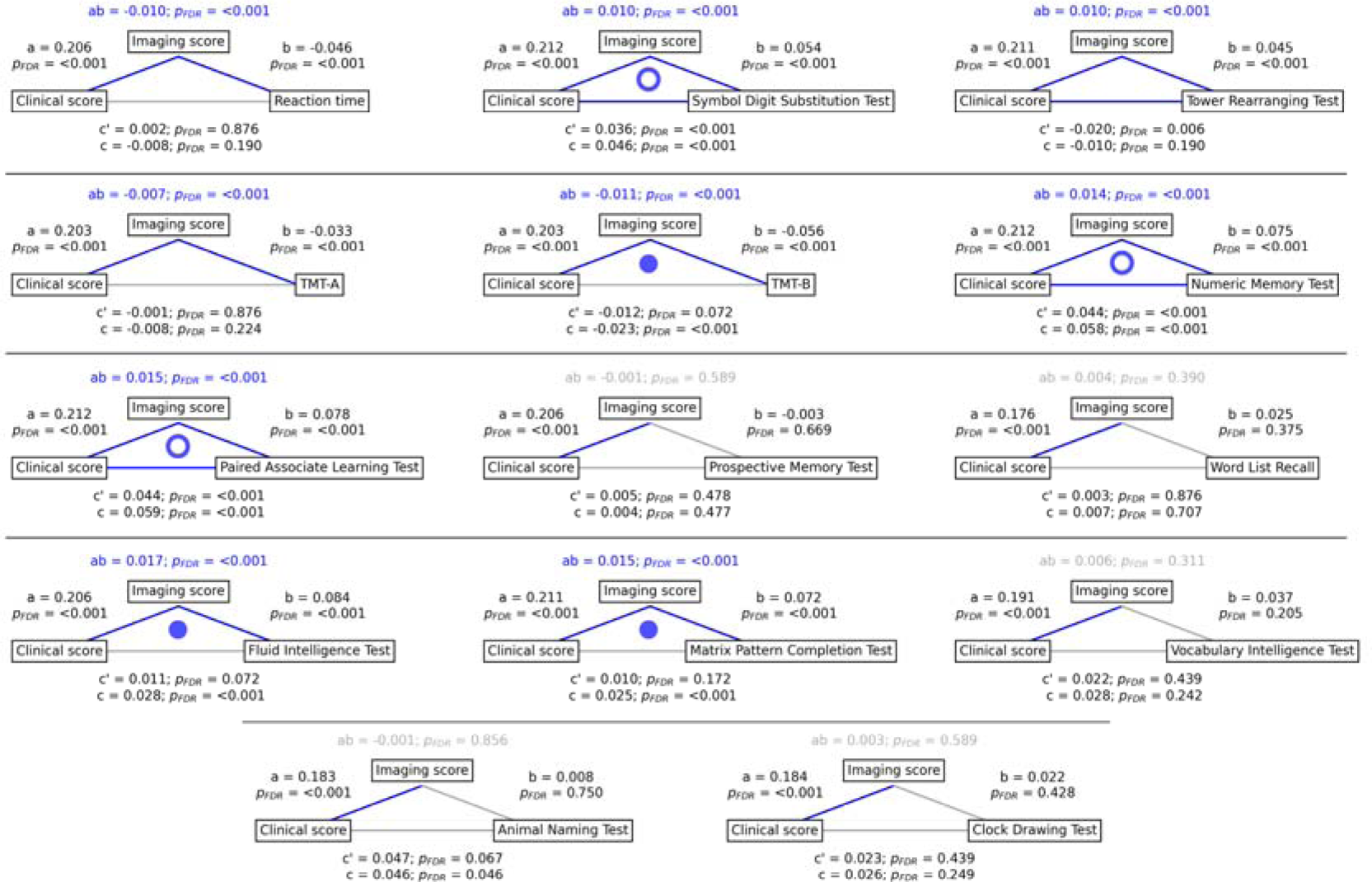
Mediation analysis results. Mediation effects of subject-specific imaging PLS scores on the relationship between MetS represented by the clinical PLS score and cognitive test performances. Path plots display standardized effects and p-values: (a) clinical score to imaging score, (b) imaging score to cognitive score, (ab) indirect effect (c’) direct effect and (c) total effect. Significant paths are highlighted in blue; non-significant in light gray. If the indirect effect ab was significant, the text for ab is highlighted in blue. A blue dot in the path plot indicates if a relationship is significantly mediated, i.e., the indirect effect ab was significant and the direct effect c’ was reduced or non-significant compared to the total effect c. An empty dot indicates a partial mediation, a full dot indicates a full mediation. *Abbreviations*: - false discovery rate-corrected p-values; PLS – partial least squares correlation; TMT-A – Trail Making Test A; TMT-B – Trail Making Test B.

### 3.4 Contextualization of MetS-associated brain morphological abnormalities

We investigated whether the pattern of MetS effects on cortical structure is conditioned by the regional density of specific cell populations and global brain network topology in a surface-based contextualization analysis (see Methods).

Therefore, we first used a virtual histology approach to relate the bootstrap ratio from PLS to the differential expression of cell-type specific genes based on microarray data from the Allen Human Brain Atlas [49]. The results are illustrated in *figure 4*. The bootstrap ratio was significantly positively correlated with the density of endothelial cells (Z_r_sp= .190, p_FDR_ = .016), microglia (Z_r_sp = .271, p_FDR_ = .016), excitatory neurons type 8 (Z_rsp_= .165, p_FDR_ = .016), inhibitory neurons type 1 (Z_r_sp = .363, p_FDR_ = .036) and excitatory neurons type 6 (Z = .146, p_FDR_ = .034) indicating that MetS-related brain morphological abnormalities are strongest in regions of the highest density of these cell types. No significant associations were found with regard to the remaining excitatory neuron types (Ex1-Ex5, Ex7), inhibitory neurons (In2-In8), astrocytes and oligodendrocytes (*supplementary table S20*). Virtual histology analysis results for bootstrap ratios corresponding with latent variables 2 and 3 are shown in *supplementary figure S21*. As a sensitivity analysis, we contextualized the t-statistic map derived from group statistics. The results remained stable except for excitatory neurons type 6 (Z_r_sp = .145, p_FDR_ = .123) and inhibitory neurons type 1 (Z_rsp_ = .432, p_FDR_ = .108), which no longer showed a significant association (*supplementary materials S22-23*).

**Figure 4.**
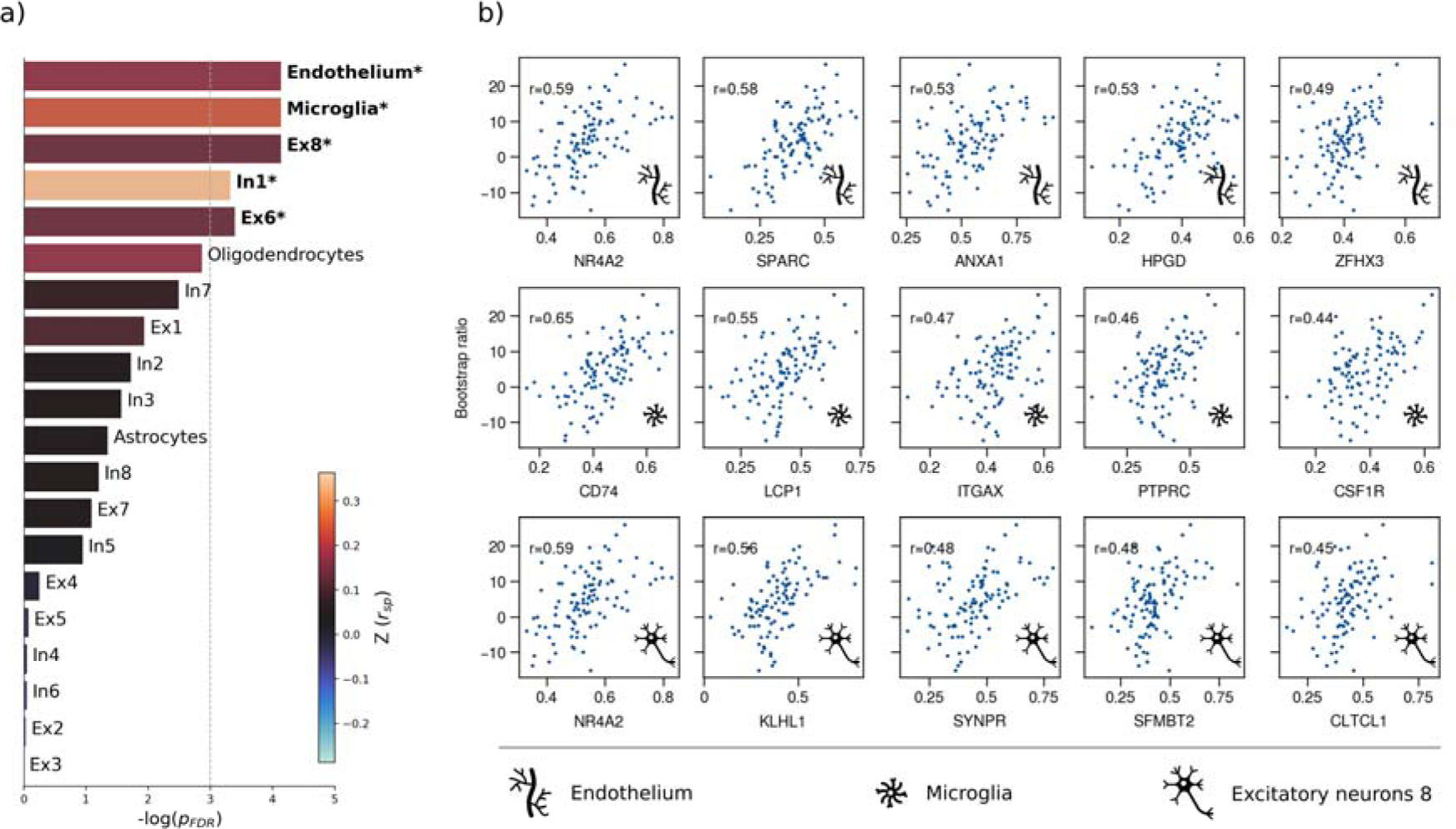
Virtual histology analysis. The correspondence between MetS effects (bootstrap ratio) and cell type-specific gene expression profiles was examined via an ensemble-based gene category enrichment analysis. a) Barplot displaying spatial correlation results. The bar height displays the significance level. Colors encode the aggregate z-transformed Spearman correlation coefficient relating the Schaefer100-parcellated bootstrap ratio and respective cell population densities. Asterisks indicate statistical significance. The significance threshold of <.05 is highlighted by a vertical dashed line. b) Scatter plots illustrating spatial correlations between MetS effects and exemplary cortical gene expression profiles per cell population significantly associated across analyses – i.e., endothelium, microglia and excitatory neurons type 8. Top 5 genes most strongly correlating with the bootstrap ratio map were visualized for each of these cell populations. Icons in the bottom right of each scatter plot indicate the corresponding cell type. A legend explaining the icons is provided at the bottom. First row: endothelium; second row: microglia; third row: excitatory neurons type 8. Virtual histology analysis results for the bootstrap ratios of latent variables 2 and 3 are shown in *supplementary figure S21*. A corresponding plot illustrating the contextualization of the t-statistic derived from group statistics is shown in *supplementary figure S22*. Abbreviations: −log(p_FDR_) – negative logarithm of the false discovery rate-corrected p-value derived from spatial lag models [36,41]; r – Spearman correlation coeffient. Z(r_sp_) – aggregate z- transformed Spearman correlation coefficient.

Second, we associated the bootstrap ratio with three pre-selected measures of brain network topology derived from group consensus functional and structural connectomes of the Human Connectome Project (HCP) (*figure 5*): weighted degree centrality (marking brain network hubs), neighborhood abnormality and macroscale functional connectivity gradients [33]. The bootstrap ratio showed a medium positive correlation with the functional neighborhood abnormality (r_sp_ = .464, p_spin_ < .001, p_smash_ < .001, p_rewire_ < .001) and a strong positive correlation with the structural neighborhood abnormality (r_sp_ = .764, p_spin_ = <.001, p_smash_ < .001, p_rewire_ < .001) indicating functional and structural interconnectedness of areas exhibiting similar MetS effects. These results remained significant when the t-statistic map was contextualized instead of the bootstrap ratio as well as when neighborhood abnormality measures were derived from consensus connectomes of the HCHS instead of the HCP (*supplementary figure S24-25*). We found no significant associations for the remaining indices of network topology, i.e., functional degree centrality (r_sp_ = .163, p_spin_ = .365, p_smash_ = .406, p_rewire_ = .870), structural degree centrality (r_sp_ = .029, p_spin_ = .423, p_smash_ = .814, p_rewire_ = .103) as well as functional cortical gradient 1 (r_sp_ = .152, p_spin_ = .313, p_smash_ = .406, p_rewire_ = .030) and gradient 2 (r_sp_ = -.177, p_spin_ = .313, p_smash_ = .406, p_rewire_ < .001).

**Figure 5.**
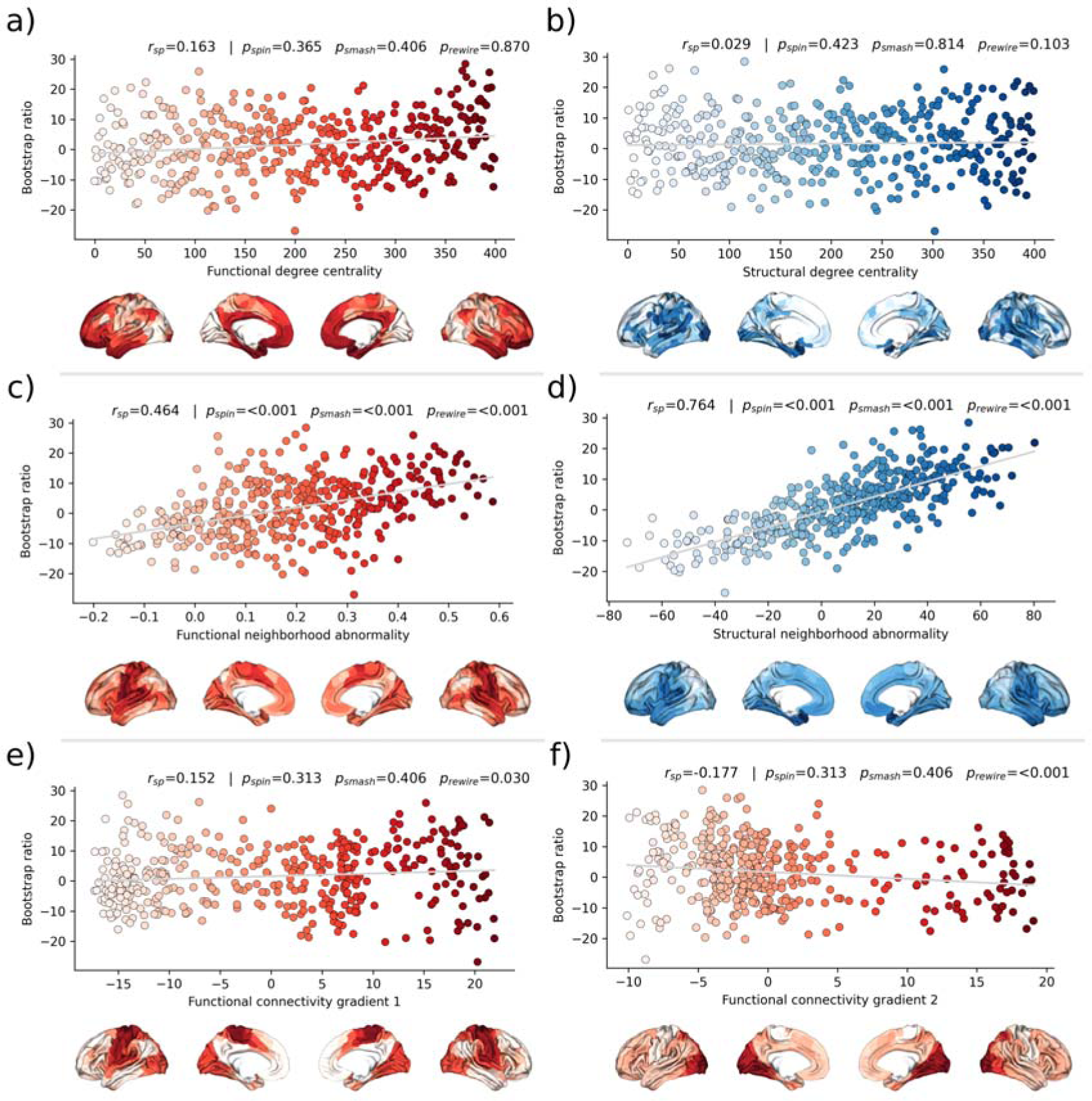
Brain network contextualization. Spatial correlation results derived from relating Schaefer400×7-parcellated maps of MetS effects (bootstrap ratio) to network topological indices (red: functional connectivity, blue: structural connectivity). Scatter plots that illustrate the spatial relationship are supplemented by respective surface plots for anatomical localization. The color coding of cortical regions and associated dots corresponds. a) & b) Functional and structural degree centrality rank. c) & d) Functional and structural neighborhood abnormality. e) & f) Intrinsic functional network hierarchy represented by functional connectivity gradients 1 and 2. Complementary results concerning t-statistic maps derived from group comparisons between MetS subjects and controls are presented in *supplementary figure S24*. Corresponding results after reperforming the analysis with HCHS derived group-consensus connectomes are presented in *supplementary figure S25.* Abbreviations: HCHS – Hamburg City Health Study; p_rewire_ - p-value derived from network rewiring [47]; p_smash_ - p-value derived from brainSMASH surrogates [46]; p_spin_ - p-value derived from spin permutation results [45]; r_sp_ - Spearman correlation coefficient.

## 4 Discussion

We investigated the impact of MetS on brain morphology and cognitive function in a large sample of individuals from two population-based neuroimaging studies. We report three main findings: 1) multivariate, data-driven statistics revealed a latent variable relating MetS and brain health: participants were distributed along a clinical-anatomical dimension of interindividual variability, linking more severe MetS to widespread brain morphological abnormalities. Negative MetS-related brain morphological abnormalities were strongest in orbitofrontal, lateral prefrontal, insular, cingulate and temporal cortices as well as subcortical areas. Positive MetS-related brain morphological abnormalities were strongest in superior frontal, parietal and occipital regions. 2) The severity of MetS was associated with executive function and processing speed, memory, and reasoning test performances, and was found to be statistically mediated by MetS-related brain morphological abnormalities. 3) The pattern of MetS-related brain morphological abnormalities appeared to be linked to regional cell composition as well as functional and structural connectivity. These findings were robust across sensitivity analyses. In sum, our study provides an in-depth examination of the intricate relationship between MetS, brain morphology and cognition.

### 4.1 PLS reveals a latent clinical-anatomical dimension relating MetS and brain health

MetS adversely impacts brain health through complex, interacting effects on the cerebral vasculature and parenchyma as shown by histopathological and imaging studies [19]. The pathophysiology of MetS involves atherosclerosis, which affects blood supply and triggers inflammation [50,51]; endothelial dysfunction reducing cerebral vasoreactivity [52]; breakdown of the blood-brain barrier inciting an inflammatory response [53]; oxidative stress causing neuronal and mitochondrial dysfunction [54]; and small vessel injury leading to various pathologies including white matter damage, microinfarcts and cerebral microbleeds [55].

To address these interacting effects, we harnessed multivariate, data-driven statistics in form of a PLS in two large-scale population-based studies to probe for covariance profiles relating the full range of MetS components (such as obesity or arterial hypertension) to regional brain morphological information in a single analysis. PLS identified seven significant latent variables with the first variable explaining the majority (71.20%) of shared variance within the imaging and clinical data (*figure 2a*). This finding indicates a relatively uniform connection between MetS and brain morphology, implying that the associative effects of various MetS components on brain structure are comparatively similar, despite the distinct pathomechanisms each component entails.

PLS revealed that all MetS components were contributing to this latent signature. However, waist circumference, hip circumference, waist-hip ratio and body mass index consistently contributed higher than the remaining variables across conducted analyses which highlights obesity as the strongest driver of MetS-related brain morphological abnormalities.

We interpret these findings as evidence that MetS-associated conditions jointly contribute to the harmful effects on brain structure rather than affecting it in a strictly individual manner. This notion is supported by previous work in the UKB demonstrating overlapping effects of individual risk factors on cortical morphology [56]. Specifically, the first latent variable related increased severity of obesity, dyslipidemia, arterial hypertension and insulin resistance with lower thickness in orbitofrontal, lateral prefrontal, insular, cingulate and temporal cortices as well as lower volume across subcortical regions (*figure 2c and d*). This profile was consistent in separate PLS analyses of UKB and HCHS participants as well as group comparisons (*supplementary figures S12* and *S16-17*). Previous research aligns with our detection of a MetS-associated frontotemporal morphometric abnormality pattern [9,13,57]. As a speculative causative pathway, human and animal studies have related the orbitofrontal, insular and anterior cingulate cortex to food-related reward processing, taste and impulse regulation [58,59]. Conceivably, structural alterations of these brain regions are linked to brain functions and behaviors that exacerbate the risk profile leading to MetS [60,61]. We also noted a positive MetS-cortical thickness association in superior frontal, parietal and occipital lobes, a less intuitive finding that has been previously reported [62,63]. Although speculative, the positive effects might be due to MetS compensating cholesterol disruptions associated with neurodegenerative processes [64].

The second latent variable accounted for 22.33% of shared variance and linked higher markers of insulin resistance and lower dyslipidemia to lower thickness and volume in lateral frontal, posterior temporal, parietal and occipital regions. The distinct covariance profile of this latent variable, compared to the first, likely indicates a separate pathomechanistic connection between MetS components and brain morphology. Given that HbA1c and blood glucose were the most significant contributors to this variable, insulin resistance might drive the observed clinical-anatomical relationship.

### 4.2 Brain morphological abnormalities mediate the relationship between MetS and cognitive deficits

Cognitive performance has been consistently linked to cardiometabolic risk factors in health and disease [65]. Yet, the pathomechanistic correlates of this relationship remain to be understood. Our mediation analysis revealed that increased MetS severity correlates with worse performance in executive function and processing speed (Symbol Digit Substitution Test, Trail Making Test B), memory (Numeric Memory Test, Paired Associate Learning Test), and reasoning (Fluid intelligence, Matrix Pattern Completion Test), with brain morphological abnormalities statistically mediating these relationships. Additionally, group comparisons indicated poorer cognitive performance in MetS subjects (*supplementary tables S9-10*) and including cognitive outcomes in the PLS as clinical variables revealed a significant contribution to the first latent variable (*supplementary materials S16b and S17b*). These results suggest that MetS is significantly associated with cognitive deficits across various domains, and brain morphological abnormalities are a crucial pathomechanistic link in this relationship. In support of this, previous studies have shown that brain structure mediates the relationship between MetS and cognitive performance in a pediatric sample and elderly patients with vascular cognitive impairment [66–68]. The detected latent variable might represent a continuous disease spectrum spanning from minor cognitive deficits due to a cardiometabolic risk profile to severe cognitive deficits due to dementia. In support of this hypothesis, the determined brain morphological abnormality pattern is consistent with the atrophy pattern found in vascular mild cognitive impairment, vascular dementia and Alzheimer’s dementia [67–69].

Collectively, these findings highlight the role of MetS in cognitive impairment and underscore the potential impact of therapies targeting cardiometabolic risk factors. Although the definitive role of such therapies in preventing cognitive decline is not yet fully established, emerging evidence suggests that these interventions can mitigate the adverse cognitive effects of MetS [70–72]. As our results highlight obesity as a key factor in the observed clinical-anatomical relationship, we think that future studies should further investigate weight-reducing interventions to examine their effects on cognitive outcomes. Advanced neuroimaging techniques promise to refine these therapeutic approaches by enabling to identify MetS patients at risk of cognitive decline that would benefit the most from targeted interventions for cognitive health protection.

### 4.3 MetS-related brain morphological abnormalities link to cellular tissue composition and network topology

To better understand the emergence of the spatial pattern of MetS-related brain morphological abnormalities, we conducted two contextualization analyses leveraging reference datasets of local gene expression data as well as properties of brain network topology.

Using a virtual histology approach based on regional gene expression data, we investigated MetS effects in relation to cell population densities (*figure 4*). As the main finding, we report that higher MetS-related brain morphological abnormalities coincide with a higher regional density of endothelial cells. This aligns with the known role of endothelial dysfunction in MetS compromising tissues via chronic vascular inflammation, increased thrombosis risk and hypoperfusion due to altered vasoreactivity and vascular remodeling [52]. As endothelial density also indicates the degree of general tissue vascularization, well-vascularized regions are also likely more exposed to cardiometabolic risk factor effects in general [50]. Our results furthermore indicate that microglial density determines a brain region’s susceptibility to MetS effects. Microglia are resident macrophages of the central nervous system sustaining neuronal integrity by maintaining a healthy microenvironment. Animal studies have linked microglial activation mediated by blood-brain barrier leakage and systemic inflammation to cardiometabolic risk [73,74]. Activated microglia can harm the brain structure by releasing reactive oxygen species, proinflammatory cytokines and proteinases [75]. Lastly, we found an association with the density of excitatory neurons of subtype 8. These neurons reside in cortical layer 6 and their axons mainly entertain long-range cortico-cortical and cortico-thalamic connections [37,76]. Consequently, layer 6 neurons might be particularly susceptible to MetS effects due to their exposition to MetS-related white matter disease [24,77]. Taken together, the virtual histology analysis indicates that MetS-related brain morphological abnormalities are associated to local cellular fingerprints. Our findings emphasize the involvement of endothelial cells and microglia in brain structural abnormalities due to cardiometabolic risk, marking them as potential targets for therapies aimed at mitigating MetS effects on brain health.

For the second approach, we contextualized MetS-related brain morphological abnormalities using principal topological properties of functional and structural brain networks. We found that regional MetS effects and those of functionally and structurally connected neighbors were correlated (*figure 5c and 5d*) – i.e., areas with similar MetS effects tended to be disproportionately interconnected. Put differently, MetS effects coincided within functional and structural brain networks. Therefore, our findings can be interpreted as evidence that a region’s functional and structural network embedding – i.e., its individual profile of functional interactions as well as white matter fiber tract connections – are associated to its susceptibility to morphological MetS effects. Multiple mechanisms might explain how connectivity might be associated to MetS-related morphological alterations. For example, microvascular pathology might impair white matter fiber tracts leading to joint degeneration in interconnected cortical brain areas: that is, the occurrence of shared MetS effects within functionally and structurally connected neighborhoods is explained by their shared (dis-)connectivity profile [78]. In support of this, previous work using diffusion tensor imaging suggests that MetS-related microstructural white matter alterations preferentially occur in the frontal and temporal lobe, which spatially matches the frontotemporal morphometric differences observed in our work [79]. Furthermore, we speculate on an interplay between local and network-topological susceptibility in MetS: functional and structural connectivity may provide a scaffold for propagating MetS-related perturbation across the network in the sense of a spreading phenomenon – i.e., a region might be influenced by network-driven exposure to regions with higher local susceptibility. Observed degeneration of a region might be aggravated by malfunctional communication to other vulnerable regions including mechanisms of excitotoxicity, diminished excitation and metabolic stress [80]. These findings underscore the relevance of brain network organization in understanding the pathomechanistic link of MetS and brain morphology.

While this work’s strengths lie in a large sample size, high-quality MRI and clinical data, robust image processing, and a comprehensive methodology for examining the link of MetS and brain health, it also has limitations. First, the virtual histology analysis relies on post-mortem brain samples, potentially different from in-vivo profiles. In addition, the predominance of UKB subjects may bias the results, and potential reliability issues of the cognitive assessment in the UKB need to be acknowledged [81]. Lastly, the cross-sectional design restricts the ability for demonstrating causative effects. Longitudinal assessment of the surveyed relationships would provide more robust evidence and therefore, future studies should move in this direction.

## 5 Conclusion

Our analysis revealed associative effects of MetS, structural brain integrity and cognition, complementing existing efforts to motivate and inform strategies for cardiometabolic risk reduction. In conjunction, a characteristic and reproducible structural imaging fingerprint associated with MetS was identified. This pattern of MetS-related brain morphological abnormalities was linked to local histological as well as global network topological features. Collectively, our results highlight how an integrative, multi-modal and multi-scale analysis approach can lead to a more holistic understanding of the neural underpinnings of MetS and its risk components. As research in this field advances, leveraging neuroimaging may improve personalized cardiometabolic risk mitigation approaches.

## Supporting information

Schaefer100×7-parcellated bootstrap ratio

Schaefer400×7-parcellated MetS effect maps

fsLR_t-statistic_group_comparison

## 6 Acknowledgements

The authors wish to acknowledge all participants and staff of the UK Biobank and Hamburg City Health Study. The participating institutes and departments from the University Medical Center Hamburg-Eppendorf contributed all with individual and scaled budgets to the overall funding of the Hamburg City Health Study. The publication has been approved by the Steering Board of the Hamburg City Health Study. This research has been conducted using the UK Biobank Resource under Application Number 41655.

Each author has made a significant contribution to the manuscript and all authors read and approved its final version. We describe individual contributions to the paper using the CRediT contributor role taxonomy. Conceptualization: M.P., F.H., B.C.; Data Curation: M.P., F.H., F.N., C.M., M.S.; Formal analysis: M.P.; Funding acquisition: S.B.E., G.T., B.C.; Investigation: M.P.; Methodology: M.P.; Software: M.P., F.H.; Supervision: G.T., B.C.; Visualization: M.P.; Writing—original draft: M.P.; Writing—review & editing: M.P., F.H., F.N., C.M., M.S., L.R., B.Z., T.Z., S.K., J.G., J.F., R.T., A.O., K.P., S.B.E., G.T., B.C.

## 8 Data availability

UK Biobank data can be obtained via its standardized data access procedure (https://www.ukbiobank.ac.uk/). HCHS participant data used in this analysis is not publicly available for privacy reasons, but access can be established via request to the HCHS steering committee. The analysis code is publicly available on GitHub (https://github.com/csi-hamburg/2023_petersen_mets_brain_morphology and https://github.com/csi-hamburg/CSIframe/wiki/Structural-processing-with-CAT).

---

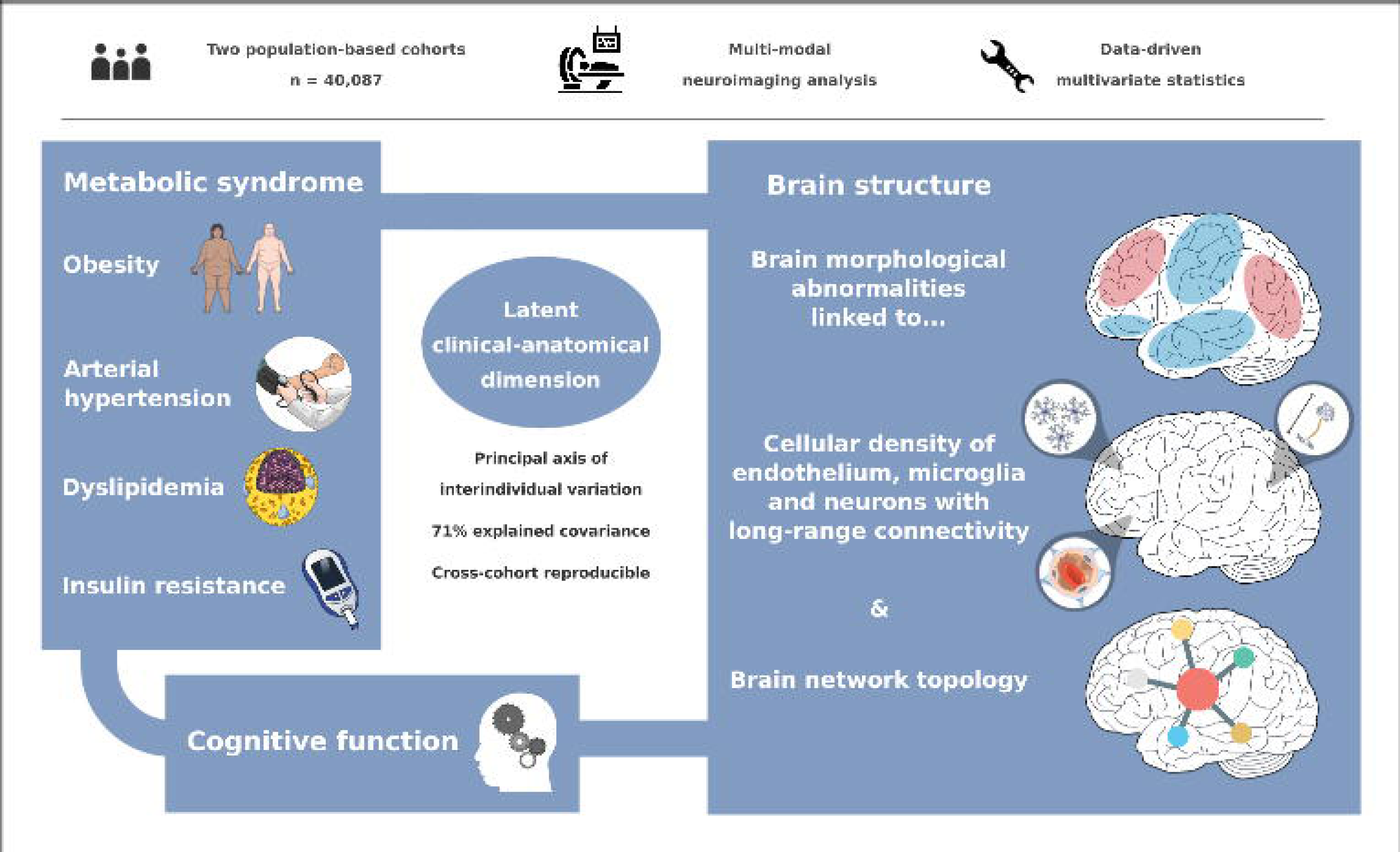

## References

[1] Alberti KGMM, Zimmet P, Shaw J. Metabolic syndrome--a new world-wide definition. A Consensus Statement from the International Diabetes Federation. Diabet Med 2006;23:469–80. 10.1111/j.1464-5491.2006.01858.x.

[2] Aguilar M, Bhuket T, Torres S, Liu B, Wong RJ. Prevalence of the metabolic syndrome in the United States, 2003-2012. JAMA 2015;313:1973–4. 10.1001/jama.2015.4260.

[3] Scuteri A, Laurent S, Cucca F, Cockcroft J, Cunha PG, Mañas LR, et al. Metabolic syndrome across Europe: different clusters of risk factors. Eur J Prev Cardiol 2015;22:486–91. 10.1177/2047487314525529.

[4] Beltrán-Sánchez H, Harhay MO, Harhay MM, McElligott S. Prevalence and trends of metabolic syndrome in the adult U.S. population, 1999-2010. J Am Coll Cardiol 2013;62:697–703. 10.1016/j.jacc.2013.05.064.

[5] Boden-Albala B, Sacco RL, Lee H-S, Grahame-Clarke C, Rundek T, Elkind MV, et al. Metabolic Syndrome and Ischemic Stroke Risk. Stroke 2008;39:30–5. 10.1161/STROKEAHA.107.496588.

[6] Atti AR, Valente S, Iodice A, Caramella I, Ferrari B, Albert U, et al. Metabolic Syndrome, Mild Cognitive Impairment, and Dementia: A Meta-Analysis of Longitudinal Studies. The American Journal of Geriatric Psychiatry 2019;27:625–37. 10.1016/j.jagp.2019.01.214.

[7] Eckel RH, Grundy SM, Zimmet PZ. The metabolic syndrome. The Lancet 2005;365:1415–28. 10.1016/S0140-6736(05)66378-7.

[8] Yates KF, Sweat V, Yau PL, Turchiano MM, Convit A. Impact of Metabolic Syndrome on Cognition and Brain: A Selected Review of the Literature. Arterioscler Thromb Vasc Biol 2012;32:2060–7. 10.1161/ATVBAHA.112.252759.

[9] Beyer F, Kharabian Masouleh S, Kratzsch J, Schroeter ML, Röhr S, Riedel-Heller SG, et al. A Metabolic Obesity Profile Is Associated With Decreased Gray Matter Volume in Cognitively Healthy Older Adults. Front Aging Neurosci 2019;11:202. 10.3389/fnagi.2019.00202.

[10] Lu R, Aziz NA, Diers K, Stöcker T, Reuter M, Breteler MMB. Insulin resistance accounts for metabolic syndromeCrelated alterations in brain structure. Hum Brain Mapp 2021;42:2434–44. 10.1002/hbm.25377.

[11] Wolf EJ, Sadeh N, Leritz EC, Logue MW, Stoop TB, McGlinchey R, et al. Posttraumatic Stress Disorder as a Catalyst for the Association Between Metabolic Syndrome and Reduced Cortical Thickness. Biological Psychiatry 2016;80:363–71. 10.1016/j.biopsych.2015.11.023.

[12] Tiehuis AM, van der Graaf Y, Mali WPTM, Vincken K, Muller M, Geerlings MI, et al. Metabolic Syndrome, Prediabetes, and Brain Abnormalities on MRI in Patients With Manifest Arterial Disease: The SMART-MR Study. Diabetes Care 2014;37:2515–21. 10.2337/dc14-0154.

[13] McIntosh EC, Jacobson A, Kemmotsu N, Pongpipat E, Green E, Haase L, et al. Does medial temporal lobe thickness mediate the association between risk factor burden and memory performance in middle-aged or older adults with metabolic syndrome? Neuroscience Letters 2017;636:225–32. 10.1016/j.neulet.2016.10.010.

[14] Sala M, de Roos A, van den Berg A, Altmann-Schneider I, Slagboom PE, Westendorp RG, et al. Microstructural Brain Tissue Damage in Metabolic Syndrome. Diabetes Care 2014;37:493–500. 10.2337/dc13-1160.

[15] Marek S, Tervo-Clemmens B, Calabro FJ, Montez DF, Kay BP, Hatoum AS, et al. Reproducible brain-wide association studies require thousands of individuals. Nature 2022:1–7. 10.1038/s41586-022-04492-9.

[16] Hamer M, Batty GD. Association of body mass index and waist-to-hip ratio with brain structure: UK Biobank study. Neurology 2019;92:e594–600. 10.1212/WNL.0000000000006879.

[17] Opel N, Thalamuthu A, Milaneschi Y, Grotegerd D, Flint C, Leenings R, et al. Brain structural abnormalities in obesity: relation to age, genetic risk, and common psychiatric disorders. Mol Psychiatry 2021;26:4839–52. 10.1038/s41380-020-0774-9.

[18] Schaare HL, Kharabian Masouleh S, Beyer F, Kumral D, Uhlig M, Reinelt JD, et al. Association of peripheral blood pressure with gray matter volume in 19- to 40-year-old adults. Neurology 2019;92:e758–73. 10.1212/WNL.0000000000006947.

[19] Borshchev YYu, Uspensky YP, Galagudza MM. Pathogenetic pathways of cognitive dysfunction and dementia in metabolic syndrome. Life Sciences 2019;237:116932. 10.1016/j.lfs.2019.116932.

[20] Fornito A, Zalesky A, Breakspear M. The connectomics of brain disorders. Nature Reviews Neuroscience 2015;16:159–72. 10.1038/nrn3901.

[21] Seidlitz J, Nadig A, Liu S, Bethlehem RAI, Vértes PE, Morgan SE, et al. Transcriptomic and cellular decoding of regional brain vulnerability to neurogenetic disorders. Nat Commun 2020;11:3358. 10.1038/s41467-020-17051-5.

[22] Miller KL, Alfaro-Almagro F, Bangerter NK, Thomas DL, Yacoub E, Xu J, et al. Multimodal population brain imaging in the UK Biobank prospective epidemiological study. Nat Neurosci 2016;19:1523–36. 10.1038/nn.4393.

[23] Jagodzinski A, Koch-gromus U, Adam G, Anders S, Augustin M. Rationale and Design of the Hamburg City Health Study. European Journal of Epidemiology 2019. 10.1007/s10654-019-00577-4.

[24] Petersen M, Frey BM, Schlemm E, Mayer C, Hanning U, Engelke K, et al. Network Localisation of White Matter Damage in Cerebral Small Vessel Disease. Scientific Reports 2020;10:9210. 10.1038/s41598-020-66013-w.

[25] Sudlow C, Gallacher J, Allen N, Beral V, Burton P, Danesh J, et al. UK Biobank: An Open Access Resource for Identifying the Causes of a Wide Range of Complex Diseases of Middle and Old Age. PLoS Med 2015;12:e1001779. 10.1371/journal.pmed.1001779.

[26] Fawns-Ritchie C, Deary IJ. Reliability and validity of the UK Biobank cognitive tests. PLoS One 2020;15:e0231627. 10.1371/journal.pone.0231627.

[27] Moms JC, Heyman A, Mohs RC, Hughes JP, van Belle G, Fillenbaum G, et al. The Consortium to Establish a Registry for Alzheimer’s Disease (CERAD). Part I. Clinical and neuropsychological assesment of Alzheimer’s disease. Neurology 1989;39:1159– 1159. 10.1212/WNL.39.9.1159.

[28] Shulman KI. Clock-drawing: is it the ideal cognitive screening test? International Journal of Geriatric Psychiatry 2000;15:548–61. 10.1002/1099-1166(200006)15:6<548::AID-GPS242>3.0.CO;2-U.

[29] Gaser C, Dahnke R, Thompson PM, Kurth F, Luders E, Initiative ADN. CAT – A Computational Anatomy Toolbox for the Analysis of Structural MRI Data 2022:2022.06.11.495736. 10.1101/2022.06.11.495736.

[30] Tian Y, Margulies DS, Breakspear M, Zalesky A. Topographic organization of the human subcortex unveiled with functional connectivity gradients. Nat Neurosci 2020;23:1421–32. 10.1038/s41593-020-00711-6.

[31] Wagner AS, Waite LK, Wierzba M, Hoffstaedter F, Waite AQ, Poldrack B, et al. FAIRly big: A framework for computationally reproducible processing of large-scale data. Sci Data 2022;9:80. 10.1038/s41597-022-01163-2.

[32] Krishnan A, Williams LJ, McIntosh AR, Abdi H. Partial Least Squares (PLS) methods for neuroimaging: A tutorial and review. NeuroImage 2011;56:455–75. 10.1016/j.neuroimage.2010.07.034.

[33] Petersen M, Nägele FL, Mayer C, Schell M, Rimmele DL, Petersen E, et al. Brain network architecture constrains age-related cortical thinning. NeuroImage 2022;264:119721. 10.1016/j.neuroimage.2022.119721.

[34] Schaefer A, Kong R, Gordon EM, Laumann TO, Zuo X-N, Holmes AJ, et al. Local-Global Parcellation of the Human Cerebral Cortex from Intrinsic Functional Connectivity MRI. Cerebral Cortex 2018;28:3095–114. 10.1093/cercor/bhx179.

[35] Lotter L. LeonDLotter/ABAnnotate: 0.1.1 2022. 10.5281/zenodo.6640855.

[36] Dukart J, Holiga S, Rullmann M, Lanzenberger R, Hawkins PCT, Mehta MA, et al. JuSpace: A tool for spatial correlation analyses of magnetic resonance imaging data with nuclear imaging derived neurotransmitter maps. Hum Brain Mapp 2021;42:555–66. 10.1002/hbm.25244.

[37] Lake BB, Ai R, Kaeser GE, Salathia NS, Yung YC, Liu R, et al. Neuronal subtypes and diversity revealed by single-nucleus RNA sequencing of the human brain. Science 2016;352:1586–90. 10.1126/science.aaf1204.

[38] Wang D, Liu S, Warrell J, Won H, Shi X, Navarro FCP, et al. Comprehensive functional genomic resource and integrative model for the human brain. Science 2018;362:eaat8464. 10.1126/science.aat8464.

[39] Markello RD, Arnatkeviciute A, Poline J-B, Fulcher BD, Fornito A, Misic B. Standardizing workflows in imaging transcriptomics with the abagen toolbox. eLife 2021;10:e72129. 10.7554/eLife.72129.

[40] Fulcher BD, Arnatkeviciute A, Fornito A. Overcoming false-positive gene-category enrichment in the analysis of spatially resolved transcriptomic brain atlas data. Nat Commun 2021;12:2669. 10.1038/s41467-021-22862-1.

[41] Burt JB, Demirtaş M, Eckner WJ, Navejar NM, Ji JL, Martin WJ, et al. Hierarchy of transcriptomic specialization across human cortex captured by structural neuroimaging topography. Nat Neurosci 2018;21:1251–9. 10.1038/s41593-018-0195-0.

[42] Lotter LD, Saberi A, Hansen JY, Misic B, Barker GJ, Bokde ALW, et al. Human cortex development is shaped by molecular and cellular brain systems 2023:2023.05.05.539537. 10.1101/2023.05.05.539537.

[43] Larivière S, Paquola C, Park B, Royer J, Wang Y, Benkarim O, et al. The ENIGMA Toolbox: multiscale neural contextualization of multisite neuroimaging datasets. Nat Methods 2021;18:698–700. 10.1038/s41592-021-01186-4.

[44] Larivière S, Rodríguez-Cruces R, Royer J, Caligiuri ME, Gambardella A, Concha L, et al. Network-based atrophy modeling in the common epilepsies: A worldwide ENIGMA study. Sci Adv 2020;6:eabc6457. 10.1126/sciadv.abc6457.

[45] Alexander-Bloch AF, Shou H, Liu S, Satterthwaite TD, Glahn DC, Shinohara RT, et al. On testing for spatial correspondence between maps of human brain structure and function. NeuroImage 2018;178:540–51. 10.1016/j.neuroimage.2018.05.070.

[46] Burt JB, Helmer M, Shinn M, Anticevic A, Murray JD. Generative modeling of brain maps with spatial autocorrelation. NeuroImage 2020;220:117038. 10.1016/j.neuroimage.2020.117038.

[47] Maslov S, Sneppen K, Zaliznyak A. Pattern Detection in Complex Networks: Correlation Profile of the Internet. Physica A: Statistical Mechanics and Its Applications 2004;333:529–40. 10.1016/j.physa.2003.06.002.

[48] Larivière S, Bayrak Ş, Vos de Wael R, Benkarim O, Herholz P, Rodriguez-Cruces R, et al. BrainStat: A toolbox for brain-wide statistics and multimodal feature associations. Neuroimage 2023;266:119807. 10.1016/j.neuroimage.2022.119807.

[49] Hawrylycz MJ, Lein ES, Guillozet-Bongaarts AL, Shen EH, Ng L, Miller JA, et al. An anatomically comprehensive atlas of the adult human brain transcriptome. Nature 2012;489:391–9. 10.1038/nature11405.

[50] Libby P, Ridker PM, Maseri A. Inflammation and atherosclerosis. Circulation 2002;105:1135–43. 10.1161/hc0902.104353.

[51] Birdsill AC, Carlsson CM, Willette AA, Okonkwo OC, Johnson SC, Xu G, et al. Low cerebral blood flow is associated with lower memory function in metabolic syndrome. Obesity 2013;21:1313–20. 10.1002/oby.20170.

[52] Lind L. Endothelium-dependent vasodilation, insulin resistance and the metabolic syndrome in an elderly cohort: The Prospective Investigation of the Vasculature in Uppsala Seniors (PIVUS) study. Atherosclerosis 2008;196:795–802. 10.1016/j.atherosclerosis.2007.01.014.

[53] Hussain B, Fang C, Chang J. Blood–Brain Barrier Breakdown: An Emerging Biomarker of Cognitive Impairment in Normal Aging and Dementia. Frontiers in Neuroscience 2021;15.

[54] Mullins CA, Gannaban RB, Khan MS, Shah H, Siddik MAB, Hegde VK, et al. Neural Underpinnings of Obesity: The Role of Oxidative Stress and Inflammation in the Brain. Antioxidants (Basel) 2020;9:1018. 10.3390/antiox9101018.

[55] Frey BM, Petersen M, Mayer C, Schulz M, Cheng B, Thomalla G. Characterization of White Matter Hyperintensities in Large-Scale MRI-Studies. Front Neurol 2019;10. 10.3389/fneur.2019.00238.

[56] Cox SR, Lyall DM, Ritchie SJ, Bastin ME, Harris MA, Buchanan CR, et al. Associations between vascular risk factors and brain MRI indices in UK Biobank. European Heart Journal 2019;40:2290–300. 10.1093/eurheartj/ehz100.

[57] Kotkowski E, Price LR, Franklin C, Salazar M, Woolsey M, DeFronzo RA, et al. A neural signature of metabolic syndrome. Hum Brain Mapp 2019:hbm.24617. 10.1002/hbm.24617.

[58] Tuulari JJ, Karlsson HK, Hirvonen J, Salminen P, Nuutila P, Nummenmaa L. Neural Circuits for Cognitive Appetite Control in Healthy and Obese Individuals: An fMRI Study. PLOS ONE 2015;10:e0116640. 10.1371/journal.pone.0116640.

[59] Rolls ET. Reward Systems in the Brain and Nutrition. Annu Rev Nutr 2016;36:435–70. 10.1146/annurev-nutr-071715-050725.

[60] Rolls ET. The orbitofrontal cortex, food reward, body weight and obesity. Social Cognitive and Affective Neuroscience 2021:nsab044. 10.1093/scan/nsab044.

[61] Price AE, Stutz SJ, Hommel JD, Anastasio NC, Cunningham KA. Anterior insula activity regulates the associated behaviors of high fat food binge intake and cue reactivity in male rats. Appetite 2019;133:231–9. 10.1016/j.appet.2018.11.011.

[62] Krishnadas R, McLean J, Batty DG, Burns H, Deans KA, Ford I, et al. Cardio-metabolic risk factors and cortical thickness in a neurologically healthy male population: Results from the psychological, social and biological determinants of ill health (pSoBid) study. NeuroImage: Clinical 2013;2:646–57. 10.1016/j.nicl.2013.04.012.

[63] Leritz EC, Salat DH, Williams VJ, Schnyer DM, Rudolph JL, Lipsitz L, et al. Thickness of the human cerebral cortex is associated with metrics of cerebrovascular health in a normative sample of community dwelling older adults. NeuroImage 2011;54:2659–71. 10.1016/j.neuroimage.2010.10.050.

[64] Qin Q, Yin Y, Xing Y, Wang X, Wang Y, Wang F, et al. Lipid Metabolism in the Development and Progression of Vascular Cognitive Impairment: A Systematic Review. Frontiers in Neurology 2021;12.

[65] Genon S, Eickhoff SB, Kharabian S. Linking interindividual variability in brain structure to behaviour. Nat Rev Neurosci 2022:1–12. 10.1038/s41583-022-00584-7.

[66] Laurent JS, Watts R, Adise S, Allgaier N, Chaarani B, Garavan H, et al. Associations Among Body Mass Index, Cortical Thickness, and Executive Function in Children. JAMA Pediatrics 2020;174:170–7. 10.1001/jamapediatrics.2019.4708.

[67] Seo SW, Ahn J, Yoon U, Im K, Lee J-M, Tae Kim S, et al. Cortical Thinning in Vascular Mild Cognitive Impairment and Vascular Dementia of Subcortical Type. Journal of Neuroimaging 2010;20:37–45. 10.1111/j.1552-6569.2008.00293.x.

[68] Kim HJ, Ye BS, Yoon CW, Noh Y, Kim GH, Cho H, et al. Cortical thickness and hippocampal shape in pure vascular mild cognitive impairment and dementia of subcortical type. Eur J Neurol 2014;21:744–51. 10.1111/ene.12376.

[69] Morys F, Potvin O, Zeighami Y, Vogel J, Lamontagne-Caron R, Duchesne S, et al. Obesity-Associated Neurodegeneration Pattern Mimics Alzheimer’s Disease in an Observational Cohort Study. J Alzheimers Dis n.d.;91:1059–71. 10.3233/JAD-220535.

[70] Veronese N, Facchini S, Stubbs B, Luchini C, Solmi M, Manzato E, et al. Weight loss is associated with improvements in cognitive function among overweight and obese people: A systematic review and meta-analysis. Neuroscience & Biobehavioral Reviews 2017;72:87–94. 10.1016/j.neubiorev.2016.11.017.

[71] Lennon MJ, Lam BCP, Lipnicki DM, Crawford JD, Peters R, Schutte AE, et al. Use of Antihypertensives, Blood Pressure, and Estimated Risk of Dementia in Late Life. JAMA Netw Open 2023;6:e2333353. 10.1001/jamanetworkopen.2023.33353.

[72] Gelber RP, Ross GW, Petrovitch H, Masaki KH, Launer LJ, White LR. Antihypertensive medication use and risk of cognitive impairment. Neurology 2013;81:888–95. 10.1212/WNL.0b013e3182a351d4.

[73] Denes A, Drake C, Stordy J, Chamberlain J, McColl BW, Gram H, et al. InterleukinC1 Mediates Neuroinflammatory Changes Associated With DietCInduced Atherosclerosis. Journal of the American Heart Association n.d.;1:e002006. 10.1161/JAHA.112.002006.

[74] Tucsek Z, Toth P, Sosnowska D, Gautam T, Mitschelen M, Koller A, et al. Obesity in Aging Exacerbates Blood–Brain Barrier Disruption, Neuroinflammation, and Oxidative Stress in the Mouse Hippocampus: Effects on Expression of Genes Involved in Beta-Amyloid Generation and Alzheimer’s Disease. The Journals of Gerontology: Series A 2014;69:1212–26. 10.1093/gerona/glt177.

[75] Dheen ST, Kaur C, Ling E-A. Microglial activation and its implications in the brain diseases. Curr Med Chem 2007;14:1189–97. 10.2174/092986707780597961.

[76] Thomson A. Neocortical layer 6, a review. Frontiers in Neuroanatomy 2010;4.

[77] Petersen M, Frey BM, Mayer C, Kühn S, Gallinat J, Hanning U, et al. Fixel based analysis of white matter alterations in early stage cerebral small vessel disease. Sci Rep 2022;12:1581. 10.1038/s41598-022-05665-2.

[78] Mayer C, Frey BM, Schlemm E, Petersen M, Engelke K, Hanning U, et al. Linking cortical atrophy to white matter hyperintensities of presumed vascular origin: Journal of Cerebral Blood Flow & Metabolism 2020. 10.1177/0271678x20974170.

[79] Segura B, Jurado MA, Freixenet N, Falcón C, Junqué C, Arboix A. Microstructural white matter changes in metabolic syndrome: a diffusion tensor imaging study. Neurology 2009;73:438–44. 10.1212/WNL.0b013e3181b163cd.

[80] Saxena S, Caroni P. Selective Neuronal Vulnerability in Neurodegenerative Diseases: from Stressor Thresholds to Degeneration. Neuron 2011;71:35–48. 10.1016/j.neuron.2011.06.031.

[81] Gell M, Eickhoff SB, Omidvarnia A, Küppers V, Patil KR, Satterthwaite TD, et al. The Burden of Reliability: How Measurement Noise Limits Brain-Behaviour Predictions 2023:2023.02.09.527898. 10.1101/2023.02.09.527898.

[82] Zeighami Y, Fereshtehnejad S-M, Dadar M, Collins DL, Postuma RB, Mišić B, et al. A clinical-anatomical signature of Parkinson’s disease identified with partial least squares and magnetic resonance imaging. NeuroImage 2019;190:69–78. 10.1016/j.neuroimage.2017.12.050.

