## Supplementary materials for "A latent clinical-anatomical dimension relating metabolic syndrome to brain structure and cognition"

#### Evidence from a Multivariate Imaging Analysis of 40,087 Individuals

Marvin Petersen[1], Felix Hoffstaedter[2,3], Felix L. Nägele[1], Carola Mayer[1], Maximilian Schell[1], D. Leander Rimmele[1], Birgit-Christiane Zyriax[4], Tanja Zeller[5], Simone Kühn[6], Jürgen Gallinat[6], Jens Fiehler[7], Raphael Twerenbold[5,8,9,10], Amir Omidvarnia[2,3], Kaustubh R. Patil[2,3], Simon B. Eickhoff[2,3], Götz Thomalla[1], Bastian Cheng[1]

[1] Department of Neurology, University Medical Center Hamburg-Eppendorf, Hamburg, Germany

[2] Institute for Systems Neuroscience, Medical Faculty, Heinrich-Heine University Düsseldorf, Düsseldorf, Germany

[3] Institute of Neuroscience and Medicine, Brain and Behaviour (INM-7), Research Center Jülich, Jülich, Germany

[4] Midwifery Science-Health Services Research and Prevention, Institute for Health Services Research in Dermatology and Nursing (IVDP), University Medical Center Hamburg-Eppendorf, 20246 Hamburg, Germany

[5] Department of General and Interventional Cardiology, University Heart and Vascular Center, Hamburg, Germany

[6] Department of Psychiatry and Psychotherapy, University Medical Center Hamburg-Eppendorf, Hamburg, Germany

[7] Department of Diagnostic and Interventional Neuroradiology, University Medical Center Hamburg-Eppendorf, Hamburg, Germany

[8] Epidemiological Study Center, University Medical Center Hamburg-Eppendorf, Hamburg, Germany

[9] German Center for Cardiovascular Research (DZHK), partner site Hamburg/Kiel/Luebeck, Hamburg, Germany

[10] University Center of Cardiovascular Science, University Heart and Vascular Center, Hamburg, Germany

### Table S1 – UK Biobank field IDs

**Table S1. UK Biobank field IDs**

| | |
|---|---|
| Age | 21003 |
| Sex | 31 |
| Education | 6133[a] |
| Waist circumference | 48 |
| Hip circumference | 49 |
| Body mass index | 21001 |
| $RR_{systolic}$ | 4080 |
| $RR_{diastolic}$ | 4079 |
| HDL | 30760 |
| LDL | 30780 |
| Cholesterol | 30690 |
| Triglycerides | 30870 |
| HbA1c | 30750 |
| Blood glucose | 30740 |
| Medication for cholesterol, blood pressure, diabetes | 6153 |
| Fluid Intelligence | 20191 |
| Matrix Pattern Completion | 6373 |
| Numeric Memory Test | 20240 |
| Paired Associate Learning | 20197 |
| Prospective Memory | 20018 |
| Reaction Time | 20023 |
| Symbol Digit Substitution | 20159 |
| Tower Rearranging Test | 21004 |
| Trail Making Test A | 6348 |
| Trail Making Test B | 6350 |

Abbreviations: RR = blood pressure
[a]Converted to International Standard Classification of Education (ISCED) via the UKBB parser (https://github.com/USC-IGC/ukbb_parser)

### Table S2 – Metabolic syndrome criteria

| **Table S2. Metabolic Syndrome Criteria of the International Diabetes Federation (IDF)**[1] | |
|---|---|
| **Metabolic syndrome = obesity + two further criteria** | |
| **Obesity** | waist circumference ♀: ≥ 80 cm; ♂: ≥ 94 cm |
| **Dyslipidemia (raised triglycerides)** | ≥ 150 mg/dL (1.7 mmol/L) or lipid lowering medication |
| **Dyslipidemia (reduced HDL cholesterol)** | ♀: < 50 mg/dL (1.29 mmol/L); ♂: < 40 mg/dL (1.03 mmol/L) in males |
| **Arterial hypertension (raised blood pressure)** | systolic BP ≥ 130 or diastolic BP ≥ 85 mm Hg<br>or antihypertensive medication or diagnosis of hypertension |
| **Insuline resistance** | Fasting plasma glucose ≥ 100 mg/dL (5.6 mmol/L)<br>or antidiabetic therapy or diagnosis of diabetes mellitus type 2[a] |

[a]Measurements of fasting plasma glucose were not available for the study sample. Consequently, the criterion of insulin resistance was only based on the diagnosis of diabetes mellitus and administration of antidiabetic therapy.

### Text S3 – Partial least squares analysis explained

Regional morphometric information (Schaefer400- and Melbourne Subcortical Atlas-parcellated) and clinical data (age sex, education and MetS component data) were arranged in two matrices $X_{n_{participants} \times n_{brain\ regions}}$ and $Y_{n_{participants} \times n_{clinical\ variables}}$ and then z-scored. Subsequently, a clinical-anatomical correlation matrix was calculated. Singular value decomposition was performed on the correlation matrix which resulted in a set of mutually orthogonal latent variables. The smaller dimension of the correlation matrix – its rank – equals the latent variable count. In our case, this was the number of clinical variables. Singular value decomposition results in a left singular vector matrix ($U_{n_{brain\ regions} \times n_{latent\ variables}}$), right singular vector matrix ($V_{n_{clinical\ variables} \times n_{latent\ variables}}$) and a diagonal matrix of singular values ($\Delta_{n_{latent\ variables} \times n_{latent\ variables}}$). Together, these represent a set of latent variables with a latent variable being composed of a left and right singular vector and a corresponding singular value. Each latent variable represents a specific covariance profile within the input data. A singular vector weights the corresponding original variables to maximize their covariance, i.e., the weighted regional values of a singular vector $U_{\text{brain regions, latent variable j}}$ can be interpreted as a maximally covarying brain morphology pattern and its corresponding clinical substrate ($V_{\text{clinical variables, latent variable j}}$). The explained variance of a latent variable was calculated as the ratio of its corresponding squared singular value to the sum of the remaining squared singular values. Significance of a latent variable was assessed by comparing the observed explained variance to a non-parametric distribution of permuted values acquired by permuting the subject order in $X$ ($n_{permute}$=5000).

Subject-specific PLS scores measure to which extent an individual expresses a covariance profile represented by a latent variable. Thus, scores can be thought of as factor weightings in factor analysis. A high score describes high agreement of a participant with the identified pattern. They were calculated by projecting $U$ on $X$ for an imaging score

$$Imaging\ score = UX$$

and $V$ on $Y$ for a clinical score

$$Clinical\ score = VY.$$

Bootstrap resampling was performed to identify brain regions and clinical variables with a high and robust contribution to the clinical-anatomical association. Individuals were randomly sampled from $X$ and $Y$ with replacement (n=5000) which resulted in a set of resampled correlation matrices propagated to singular value decomposition resulting in a sampling distribution of singular vector weights for each input variable. This enabled the computation of 95% confidence intervals for the clinical variables and a bootstrap ratio for the brain regions.

$$Bootstrap\ ratio = \frac{Singular\ vector\ weight\ U_{\text{brain region i, latent variable j}}}{Standard\ error\ estimated\ from\ bootstrapping}.$$

The bootstrap ratio measures a brain region's contribution to the observed covariance profile of a respective latent variable, as a relevant region shows a high singular vector weight alongside a small standard error implying stability across bootstraps.

### Text S4 – Cortical indices of brain network topology

Connectivity metrics were based on indices derived from group-level Schaefer400x7-parcellated functional and structural connectomes derived from the Human Connectome Young Adults dataset included in the ENIGMA toolbox.[2] The preprocessing of these connectomes is described elsewhere.[3]

Weighted degree centrality. Weighted degree centrality is a measure of a network node's (here, a network node equals a Schaefer400x7 parcel as described above) relevance and is commonly used for identification of brain network hubs.[4] The degree centrality of a node $i$ was computed as the sum of its functional or structural connection weights.[5] The resulting values were ranked before further analysis.

Neighborhood abnormality. Neighborhood abnormality represents a summary measure of a cortical property in the node neighborhood defined by functional or structural brain network connectivity.[6] In this work, the MetS-related thickness abnormalities (bootstrap ratio or t-statistic) in nodes $j$ connected to node $i$ were averaged and weighted by their respective functional or structural seed connectivity ($w_{ij}$):

$$A_i = \frac{1}{N_i} \sum_{j \in N_i} C_j \, w_{ij}$$

where $j$ is one of the connected nodes $N_i$, $C_j$ is the measure of MetS-related effects on cortical thickness and the corresponding connection weight $w_{ij}$. The term $\frac{1}{Ni}$ corrects for nodal degree by normalizing by the number of connections. For example, a high positive or negative $A_i$ represents strong connectivity to nodes of pronounced MetS effects.[7]

Functional connectivity gradients. To contextualize the MetS-related effects on cortical thickness with the functional network hierarchy, we derived macroscale functional connectivity gradients as a proxy of the canonical sensorimotor-association axis, which demonstrably determines the distribution of manifold cortical properties.[8,9] Functional connectivity gradients were derived by applying diffusion map embedding on the HCP functional connectivity matrix using BrainSpace.[10] A functional connectivity gradient can be interpreted as a spatial axis of connectivity variation spanning the cortical surface, as nodes of similar connectivity profiles are closely located on these axes.

### Figure S5 – Flowchart sample selection procedure

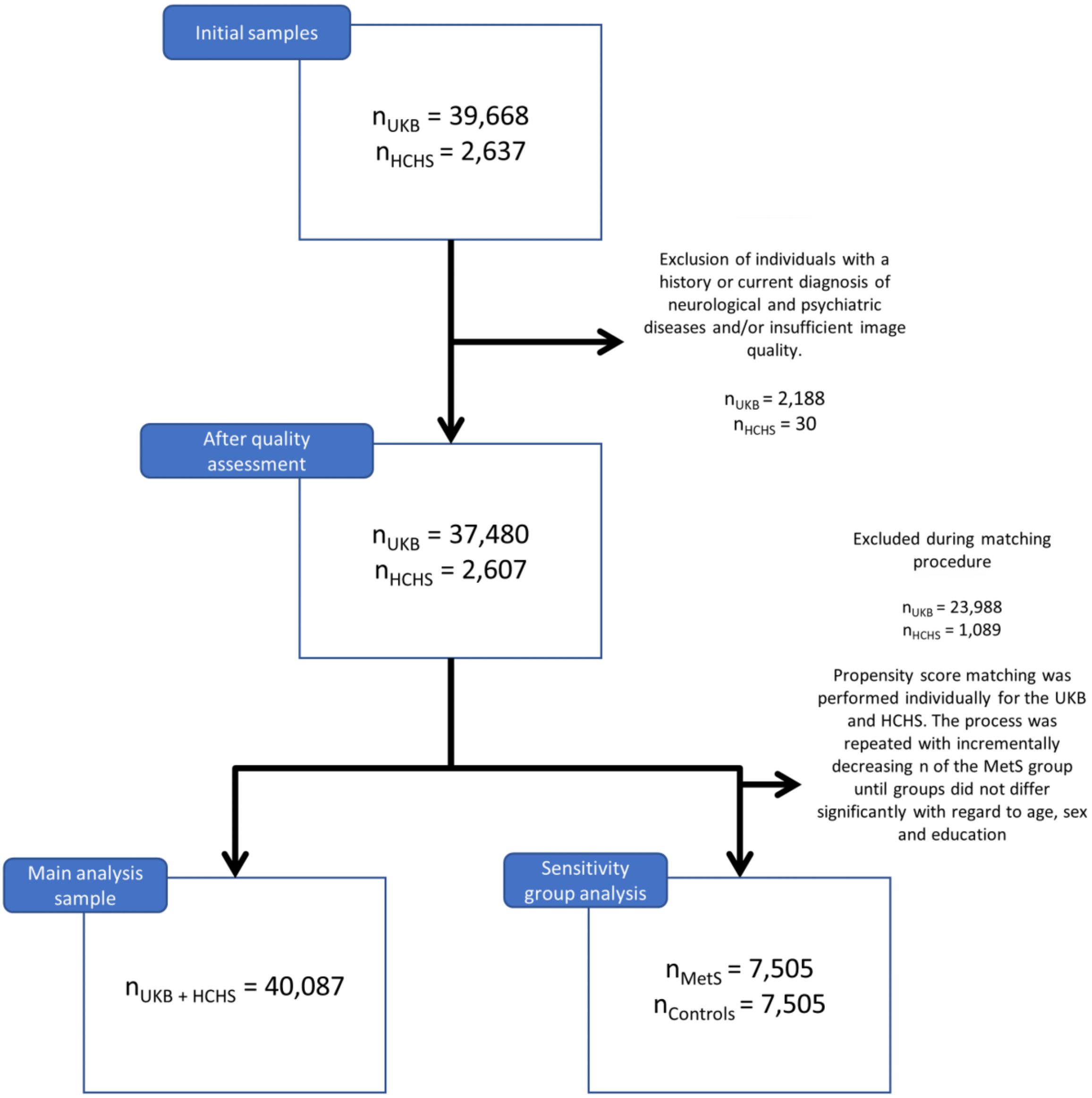

### Text S6 – Case-control analysis

As a sensitivity analysis and to facilitate the comparison with previous reports which mainly rely on group statistics, we supplemented the continuous partial least squares correlation analysis with a group analysis based on a case-control design.

Matching procedure

After quality assessment, individuals with metabolic syndrome were identified based on the consensus criteria of the *International Diabetes Federation* ($n_{UKB}$=6746, $n_{HCHS}$=759; for details on the definition see *supplementary table S2*.).[1] An individual was considered to exhibit MetS in case of obesity (increased waist circumference) and two further criteria being raised plasma triglycerides, reduced HDL cholesterol, arterial hypertension or insulin resistance. Of note, measurements of fasting plasma glucose were not available for the study sample. Consequently, the criterion of insulin resistance was only based on the diagnosis of diabetes mellitus and administration of antidiabetic therapy. Within each cohort, an equally sized control cohort was sampled which was matched for age, sex and education (International Standard Classification of Education) using propensity score matching as implemented in the matchit (v4.3.3) R package.[11] MetS and control samples from both cohorts were pooled yielding an analysis sample of 15,010 individuals ($n_{MetS}$=7505, $n_{controls}$=7505). For a flowchart providing details on the sample selection procedure please refer to *supplementary figure S5* (see above). For detailed matching results refer to supplementary figures S7-8.

Group comparison of clinical data

Sample characteristics were compared between participants with MetS and controls using $\chi^2$-tests for binary and two-sample t-tests for continuous data. Cognitive variables were compared within UKB and HCHS subgroups via analyses of covariance (ANCOVA) adjusting for age, sex and education. Resulting test statistics were converted to Cohen's d which quantifies the group difference in standard deviations. P-values were false-discovery rate (FDR)-corrected for multiple comparisons. Separate group statistics of demographic, risk and cognitive variables for the UKB and HCHS are shown in *supplementary tables S9-10*. Individuals with MetS exhibited a more severe risk profile indicating that the group definitions captured considerable differences in the MetS components profile. Group differences regarding MetS criteria proportions are visualized in *supplementary figure S11*. As the cognitive assessment of the UKB and HCHS differed, cognitive scores were compared between groups within the individual studies. Corresponding results are listed in *supplementary tables S9 and S10*. UKB subjects with MetS performed significantly worse in the Fluid Intelligence Test (6.66 ± 2.10 vs. 6.82 ± 2.09, Cohen's d = .08, $p_{FDR}$ < .001), Numeric Memory Test (6.64 ± 1.61 vs. 6.84 ± 1.53, Cohen's d = .12, $p_{FDR}$ < .001), Paired Associate Learning Test (6.45 ± 2.60 vs. 6.73 ± 2.61, Cohen's d = .10, $p_{FDR}$ < .001) and Symbol Digit Substitution Test (18.47 ± 5.12 vs. 19.00 ± 5.16, Cohen's d = .10, $p_{FDR}$ < .001). HCHS subjects exhibiting MetS showed worse cognitive performance in the Animal Naming Test (23.71 ± 6.46 vs. 24.77 ± 6.75, Cohen's d = .16, $p_{FDR}$ < .009) and Multiple-choice Vocabulary Intelligence Test (31.18 ± 3.43 vs. 31.71 ± 3.22, Cohen's d = .16, $p_{FDR}$ < .034).

Vertex-wise cortical thickness analysis

The cortical thickness of individuals with MetS and matched controls were compared on a surface vertex-level leveraging the BrainStat toolbox (v 0.3.6, https://brainstat.readthedocs.io/).[12] The corresponding results are shown in *supplementary figure S12* for the pooled group (n=15,010). The vertex-wise t-statistic, which captures the differential MetS effects across the cortical surface, was Schaefer400 and Schaefer100-parcellated and propagated to further analyses. The t-statistic map strongly correlated with the bootstrap ratio maps derived from the PLS analyses (*supplementary figure S18*). Furthermore, the t-statistic map was significantly associated with density of endothelial cells ($Z_{r_{sp}}$ = .208, $p_{FDR}$ = .040), microglia ($Z_{r_{sp}}$ = .321, $p_{FDR}$ = .040), excitatory neurons type 8 ($Z_{r_{sp}}$ = .208, $p_{FDR}$ = .004) and also correlated significantly with the functional neighborhood abnormality ($r_{sp}$ = .313, $p_{spin}$ = .024, $p_{smash}$ = .018, $p_{rewire}$ < .001) and structural neighborhood abnormality ($r_{sp}$ = .775, $p_{spin}$ = <.001, $p_{smash}$ < .001, $p_{rewire}$ < .001) (supplementary materials S22-24).

#### Figure S7 – Matching - UK Biobank

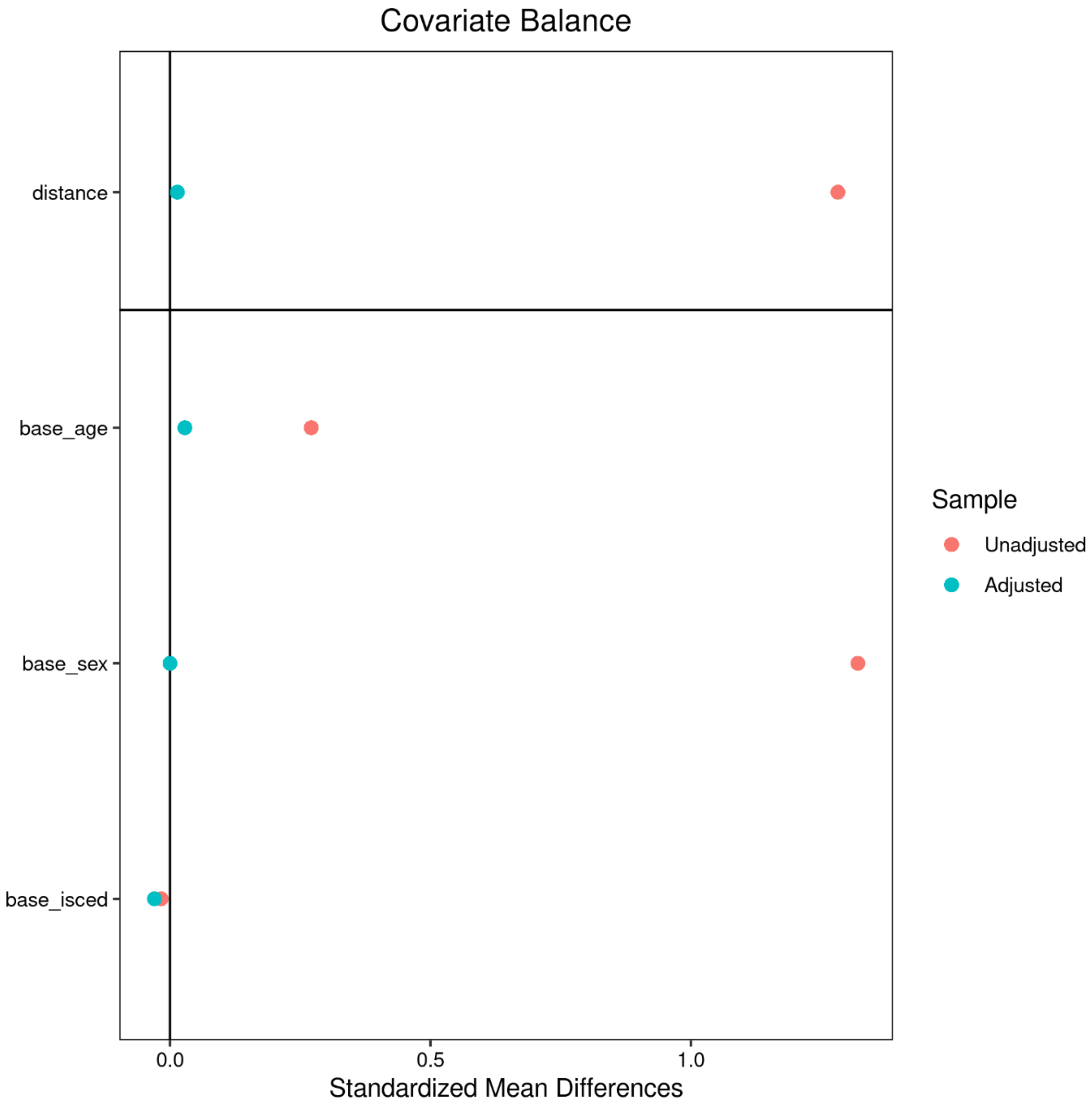

#### Figure S8 – Matching – Hamburg City Health Study

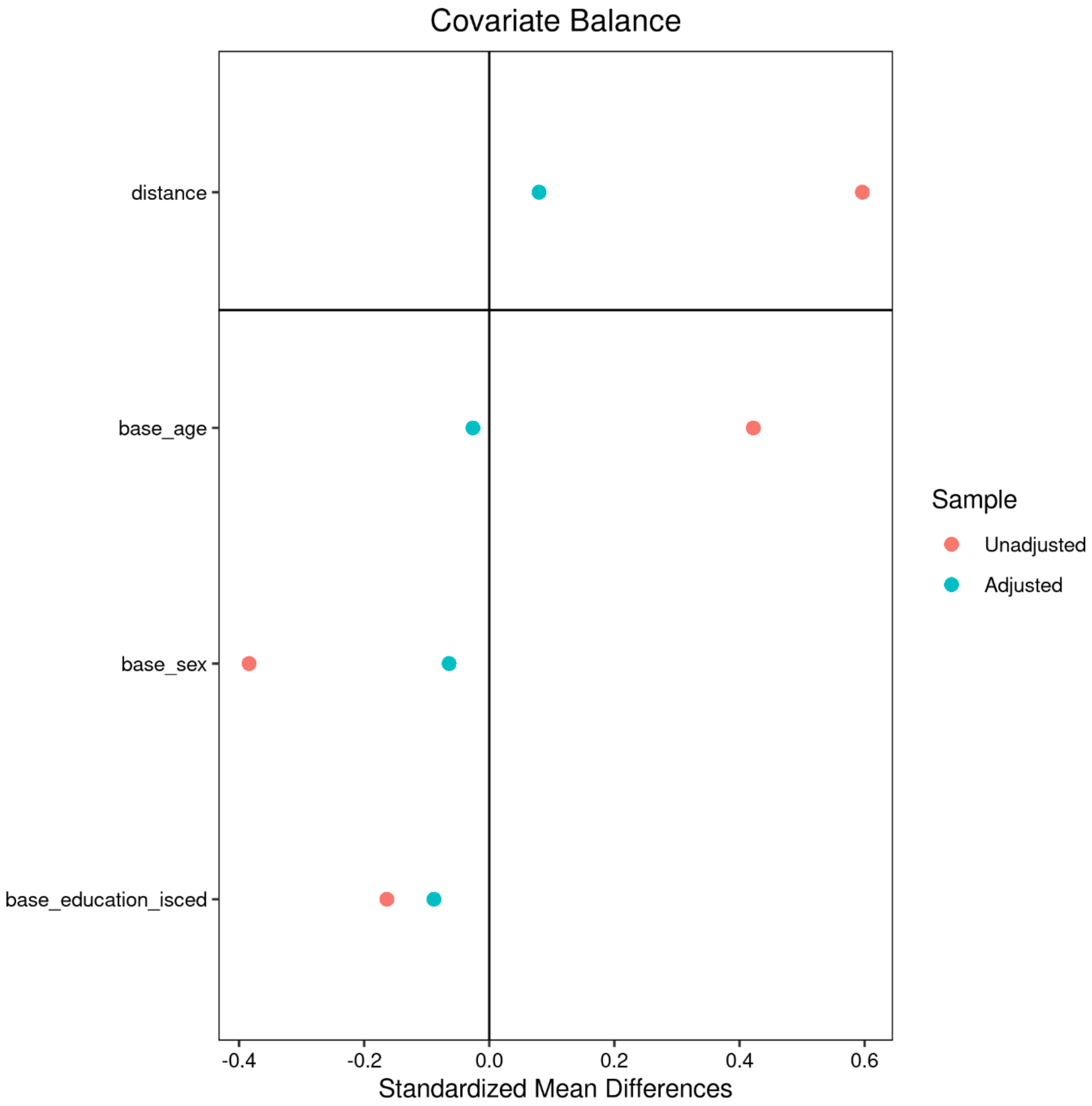

### Table S9 – Descriptive group statistics - UK Biobank

**Table S9. Descriptive group statistics UKB**

| Metric[a] | Individuals with MetS | Matched controls | $P_{uncorr}$ | $P_{FDR}$ | *Stat*[b] |
|---|---|---|---|---|---|
| **Age (years)** | 64.73 ± 7.42 (6746) | 64.51 ± 7.27 (6746) | .095 | .154 | -.03 |
| **Sex (% female)** | 18.81 (6746) | 18.81 (6746) | >.99 | >.99 | 0 |
| **Education (ISCED)** | 2.63 ± 0.73 (6746) | 2.67 ± 0.71 (6746) | .036 | .069 | .04 |
| **<u>Metabolic syndrome criteria</u>** | | | | | |
| **Waist circumference (cm)** | 97.39 ± 10.21 (6726) | 88.22 ± 10.59 (6595) | <.001 | **<.001**** | -.88 |
| **$RR_{systolic}$ (mmHg)** | 146.41 ± 15.38 (6213) | 135.57 ± 17.71 (5397) | <.001 | **<.001**** | -.66 |
| **$RR_{diastolic}$ (mmHg)** | 82.26 ± 9.39 (6214) | 77.58 ± 9.79 (5397) | <.001 | **<.001**** | -.49 |
| **Antihypertensive therapy (%)** | 9.96 (6746) | 9.68 (6746) | <.001 | **<.001**** | 7.07 |
| **HDL (mmol/L)** | 1.18 ± 0.26 (6225) | 1.49 ± 0.32 (6332) | <.001 | **<.001**** | 1.08 |
| **Triglycerides (mmol/L)** | 2.43 ± 1.13 (6617) | 1.30 ± 0.59 (6543) | <.001 | **<.001**** | -1.25 |
| **Lipid lowering therapy (%)** | 39.05 (6746) | 7.07 (6746) | <.001 | **<.001**** | 2446.5 |
| **Blood glucose (mmol/L)** | 5.18 ± 1.41 (6219) | 4.92 ± 0.68 (6325) | <.001 | **<.001**** | -.23 |
| **Antidiabetic therapy (%)** | 0.06 (6746) | 0.19 (6746) | .052 | .097 | 3.77 |
| **<u>Cognitive scores</u>** | | | | | |
| **Fluid Intelligence** | 6.66 ± 2.10 (6221) | 6.82 ± 2.09 (6241) | <.001 | **<.001**** | .08 |
| **Matrix Pattern Completion** | 8.02 ± 2.13 (4283) | 8.14 ± 2.06 (4355) | .055 | .096 | .06 |
| **Numeric Memory Test** | 6.64 ± 1.61 (4419) | 6.84 ± 1.53 (4505) | <.001 | **<.001**** | .12 |
| **Paired Associate Learning** | 6.45 ± 2.60 (4337) | 6.73 ± 2.61 (4392) | <.001 | **<.001**** | .10 |
| **Prospective Memory** | 1.05 ± 0.40 (6362) | 1.06 ± 0.39 (6349) | .221 | .339 | .02 |
| **Reaction Time** | 590.75 ± 108.27 (6331) | 590.15 ± 111.13 (6325) | .792 | .858 | -.005 |
| **Symbol Digit Substitution** | 18.47 ± 5.12 (4292) | 19.00 ± 5.16 (4353) | <.001 | **<.001**** | .10 |
| **Tower Rearranging Test** | 10.00 ± 3.28 (4255) | 10.08 ± 3.20 (4325) | .747 | .845 | .02 |
| **Trail Making Test A (sec)** | 226.83 ± 86.26 (4337) | 224.06 ± 83.06 (4392) | .643 | .836 | -.03 |
| **Trail Making Test B (sec)** | 561.81 ± 271.39 (4337) | 553.62 ± 282.55 (4392) | .611 | .756 | -.03 |

| | | | | | |
|---|---|---|---|---|---|
| **Imaging** | | | | | |
| **Mean cortical thickness (mm)** | 2.392 ± 0.09 (6746) | 2.397 ± 0.09 (6746) | .035 | .071 | .05 |

Abbreviations: cm = centimeter, dL = deciliter, HDL = high density lipoprotein, ISCED = International Standard Classification of Education, MetS = metabolic syndrome, mg = milligram, mm = millimeter, mmHg = millimeters of mercury, mmol/L = millimole per liter, PC = principal component, $P_{uncor}$ = uncorrected p-values, $P_{FDR}$ = false-discovery rate-corrected p-values, RR = Blood pressure, sec = seconds

[a]Presented as mean ± SD (N)

[b]Presented as $\chi 2$ for categorical and Cohen's d for continuous data

** Denotes statistical significance at FDR-corrected *P* <.001

### Table S10 – Descriptive group statistics - Hamburg City Health Study

**Table S10. Descriptive group statistics HCHS**

| Metric[a] | Individuals with MetS | Matched controls | $P_{uncorr}$ | $P_{FDR}$ | *Stat* |
|---|---|---|---|---|---|
| **Age (years)** | 65.77 ± 7.40 (759) | 65.97 ± 7.52 (759) | 0.613 | 0.647 | 0.03 |
| **Sex (% female)** | 33.47 | 36.50 | 0.236 | 0.281 | 1.4 |
| **Education (ISCED)** | 2.37 ± 0.58 (759) | 2.42 ± 0.60 (759) | .09 | .114 | .09 |
| **Metabolic syndrome criteria** | | | | | |
| **Waist circumference (cm)** | 103.38 ± 11.23 (754) | 91.45 ± 11.36 (747) | <.001 | **<.001**** | -1.06 |
| **$RR_{systolic}$ (mmHg)** | 145.66 ± 18.54 (740) | 140.17 ± 21.10 (746) | <.001 | **<.001**** | -.28 |
| **$RR_{diastolic}$ (mmHg)** | 83.75 ± 10.11 (740) | 82.00 ± 10.40 (746) | .001 | **.002** | -.17 |
| **Antihypertensive therapy (%)** | 52.60% | 26.22% | <.001 | **<.001**** | 108.89 |
| **HDL (mg/dL)** | 54.60 ± 16.13 (751) | 67.63 ± 17.46 (759) | <.001 | **<.001**** | .78 |
| **Triglycerides (mg/dL)** | 161.53 ± 92.61 (751) | 91.23 ± 30.62 (759) | <.001 | **<.001**** | -1.02 |
| **Lipid lowering therapy (%)** | 40.85% | 7.64% | <.001 | **<.001**** | 225.29 |
| **Blood glucose (mg/dL)** | 107.47 ± 28.59 (742) | 90.99 ± 10.87 (753) | <.001 | **<.001**** | -.76 |
| **Antidiabetic therapy (%)** | 14.42% | 1.45% | <.001 | **<.001**** | 85.47 |
| **Cognitive scores** | | | | | |
| **Animal Naming Test** | 23.71 ± 6.46 (712) | 24.77 ± 6.75 (711) | .005 | **.009** | .16 |
| **Clock Drawing Test** | 6.36 ± 1.17 (730) | 6.39 ± 1.16 (733) | .774 | .774 | .02 |
| **Trail Making Test A (sec)** | 41.26 ± 14.28 (685) | 40.42 ± 14.54 (675) | .321 | .359 | -.06 |
| **Trail Making Test B (sec)** | 93.74 ± 37.30 (675) | 89.89 ± 37.69 (671) | .086 | .114 | -.10 |
| **Multiple-Choice Vocabulary Intelligence Test** | 31.18 ± 3.43 (603) | 31.71 ± 3.22 (619) | .019 | **.034** | .16 |
| **Word List Recall** | 7.42 ± 1.89 (691) | 7.64 ± 1.84 (673) | .057 | .083 | .12 |
| **Imaging** | | | | | |
| **Mean cortical thickness (mm)** | 2.327 ± 0.08 (759) | 2.334 ± 0.08 (757) | .045 | .071 | .1 |

Abbreviations: cm = centimeter, dL = deciliter, HDL = high density lipoprotein, ISCED = International Standard Classification of Education, MetS = metabolic syndrome, mg = milligram, mm = millimeter, mmHg = millimeters of mercury, $P_{uncor}$ = uncorrected p-values, $P_{FDR}$ = false-discovery rate-corrected p-values, RR = Blood pressure, sec = seconds

[a]Presented as mean ± SD (N)
** Denotes statistical significance at FDR-corrected $P$ <.001

### Figure S11 – Proportion of metabolic syndrome criteria

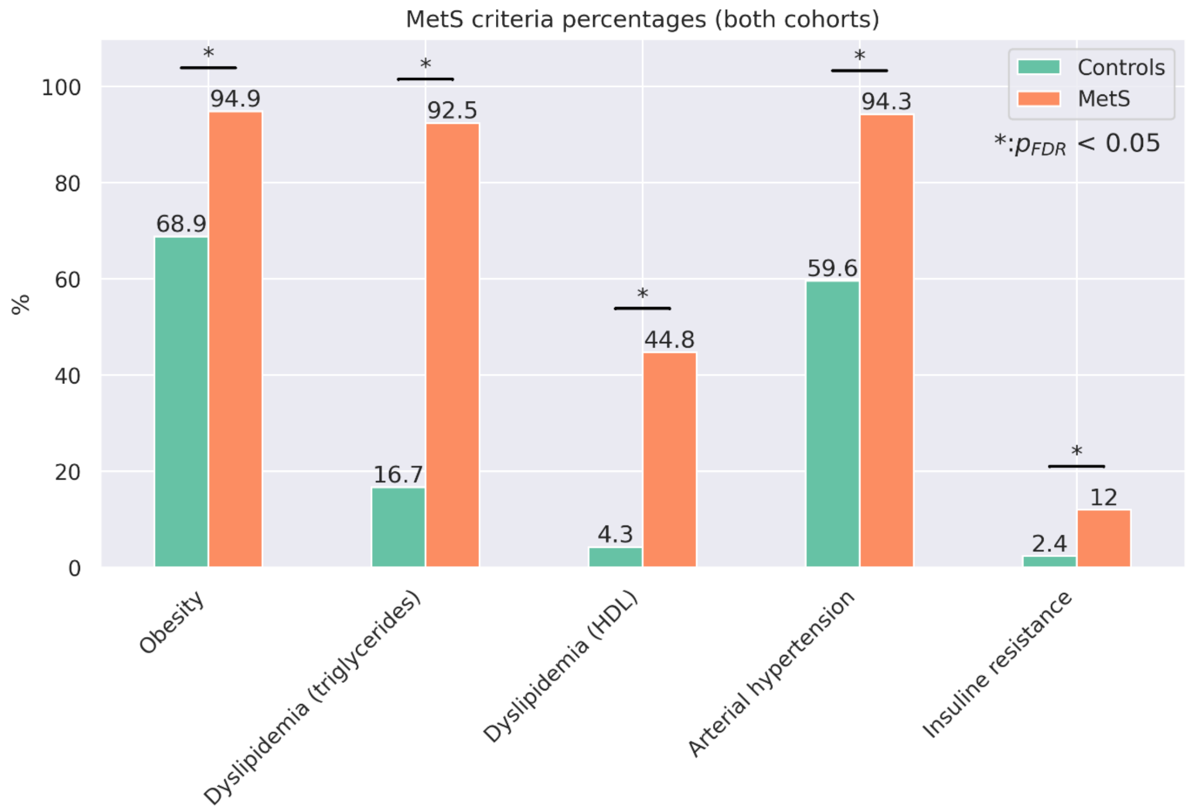

Barplots indicate the percentage amount of MetS criteria that apply by group for the pooled sample. Significant group differences in $\chi^2$-tests are highlighted by asterisks.

### Figure S12 – Vertex-wise group comparison of cortical thickness

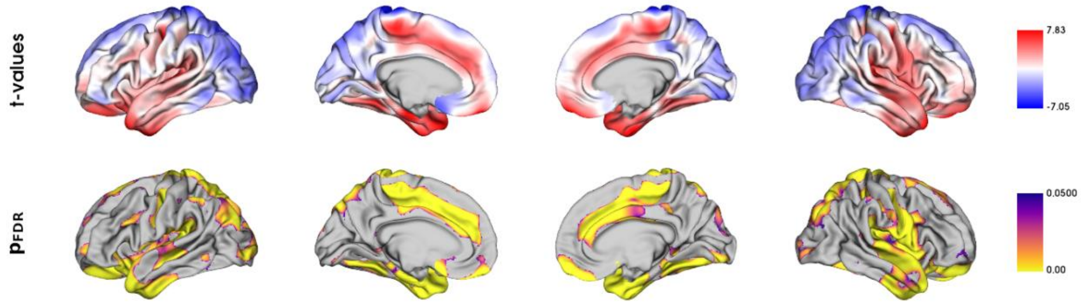

Vertex-level group comparison between individuals with metabolic syndrome and matched controls. Resulting surface maps of standardized *t*-statistic estimates encode the group-differences between patients and controls, with lower cortical thickness in the MetS group being represented by a positive *t* and lower by a negative *t*. The vertex-wise t-statistic map was Schaefer-parcellated for the downstream analyses.

### Figure S13 – Correlation matrix of metabolic syndrome-related risk factors

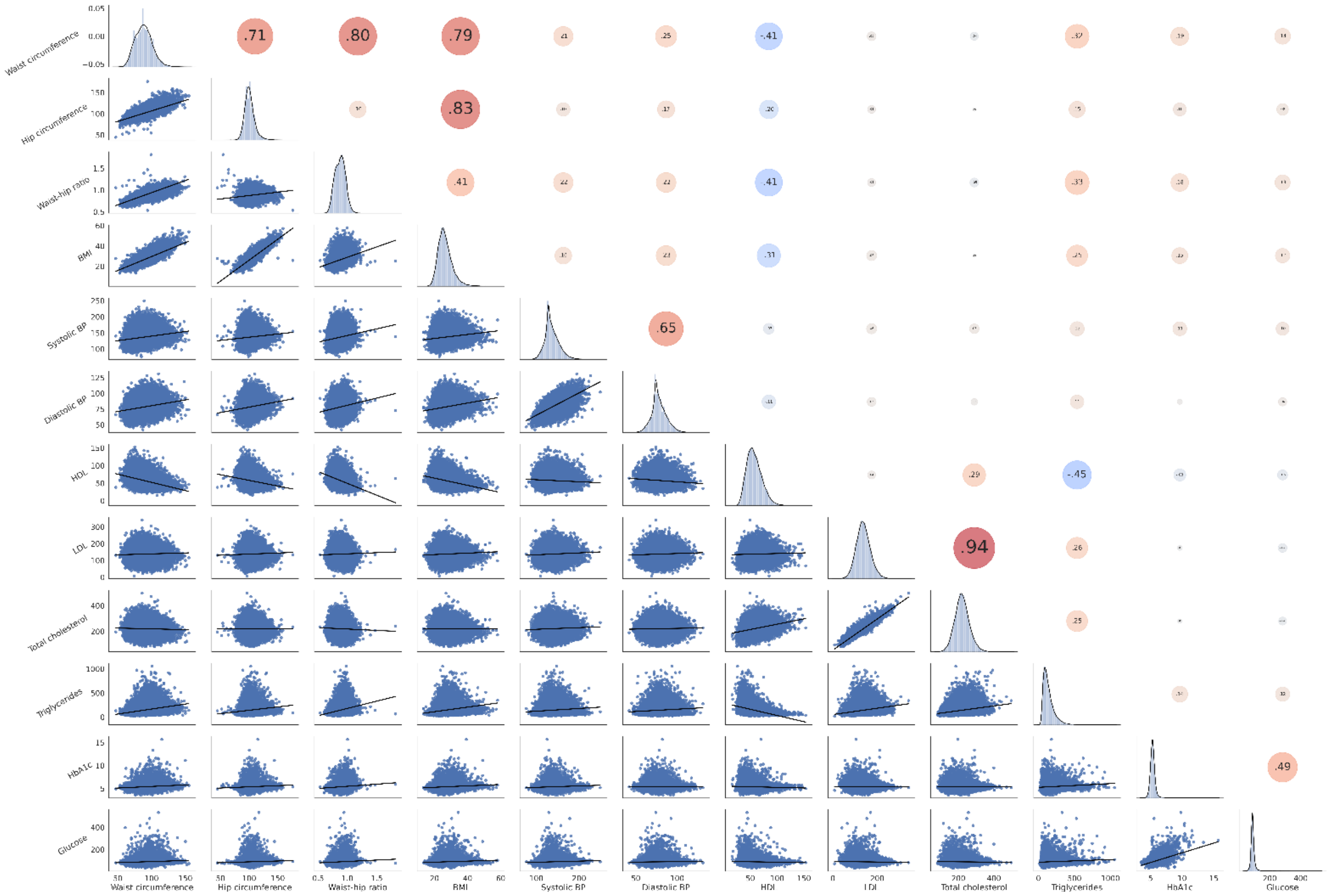

The upper triangle of the matrix displays Pearson correlations with dot size and color representing the magnitude of the coefficients. The diagonal shows kernel density plots. The lower triangle illustrates the variables' linear relationships via regression plots. Of note, non-fasting plasma glucose was investigated in this analysis. Abbreviations: PC – principal component; W. circ. – waist circumference.

### Table S14 – Partial least squares analysis – Latent variables

| Latent variable | Explained variance (%) | p-value |
|---|---|---|
| 0 | 71.20 | 0.0002 |
| 1 | 22.33 | 0.0002 |
| 2 | 2.12 | 0.0002 |
| 3 | 1.84 | 0.0006 |
| 4 | 1.03 | 0.0026 |
| 5 | 0.52 | 0.0266 |
| 6 | 0.38 | 0.0100 |
| 7 | 0.23 | 0.0032 |
| 8 | 0.18 | 0.1178 |
| 9 | 0.16 | 0.2122 |
| 10 | 0.00 | 0.3137 |
| 11 | 0.00 | 0.0608 |
| 12 | 0.00 | 1 |
| 13 | 0.00 | 1 |
| 14 | 0.00 | 1 |
| 15 | 0.00 | 1 |

#### Figure S15 – Partial least squares analysis – Latent variable 2

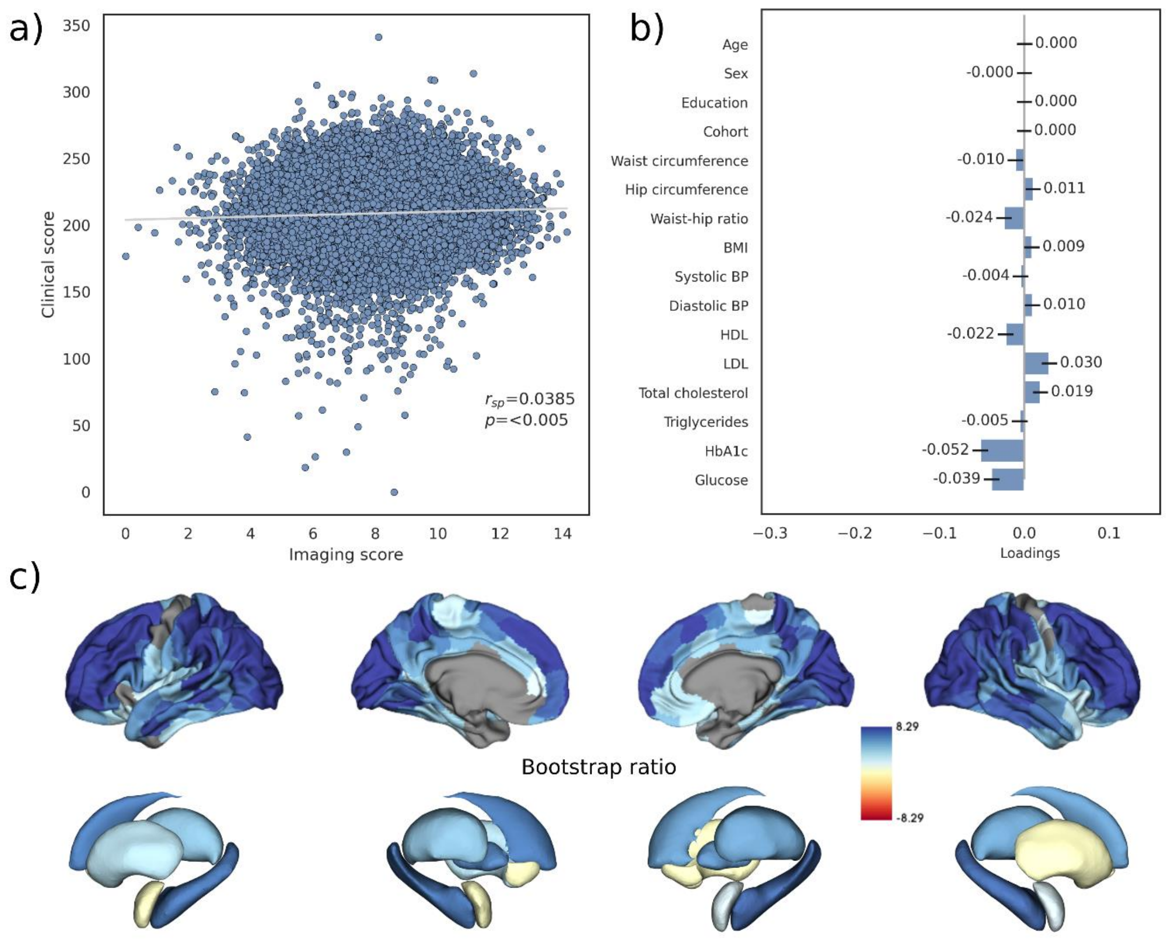

The figure presents results of latent variable 2 of the partial least squares correlation presented in the main manuscript (*figure 1*). Abbreviations: BMI – Body mass index, HDL – high density lipoprotein, LDL – low density lipoprotein.

### Figure S16 – Partial least squares analysis - UK Biobank (including cognitive test results)

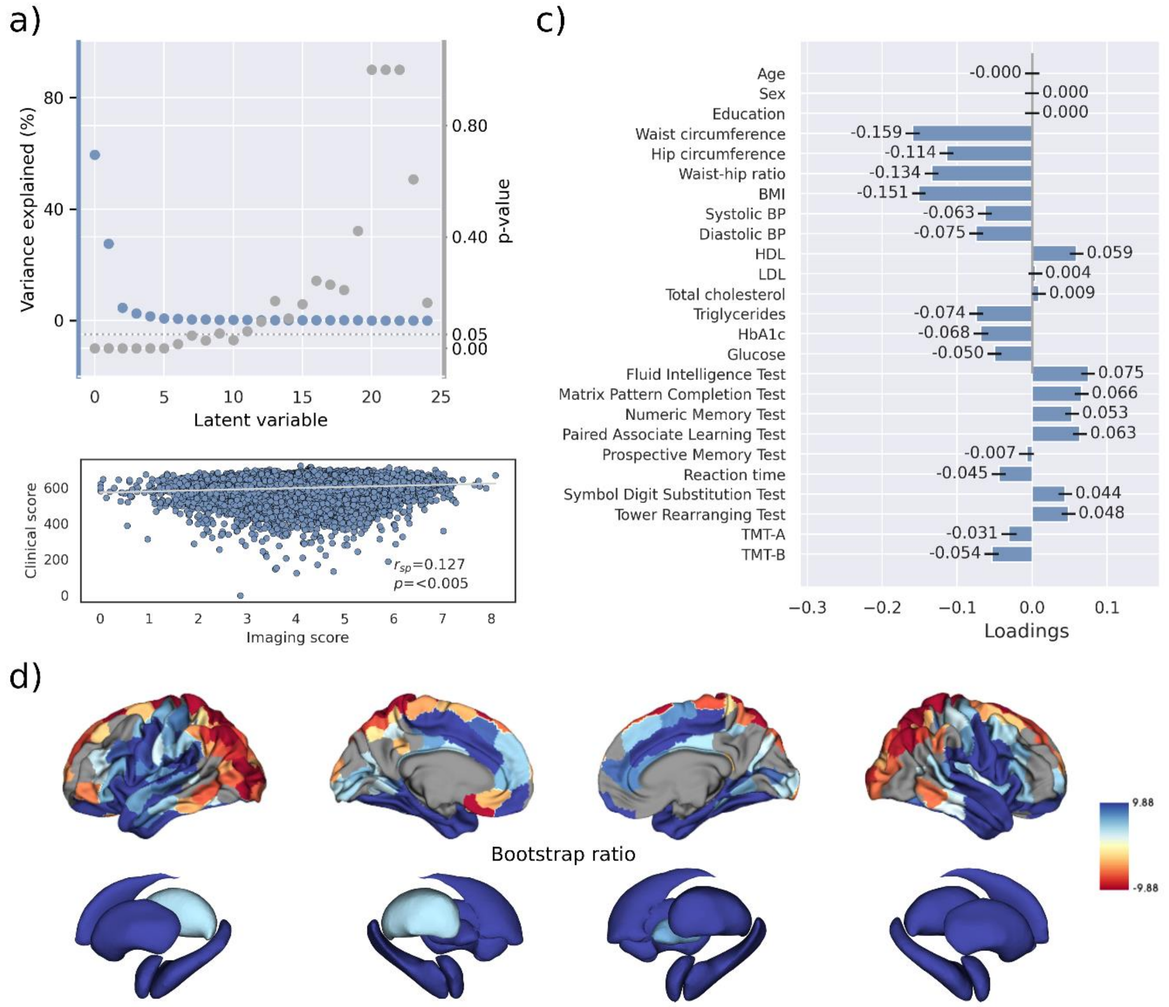

Partial least squares analysis of the UK Biobank subsample including cognitive test results. a) Explained variance and p-values of latent variables. b) Scatter plot relating subject-specific clinical and imaging scores. Higher scores indicate higher adherence to the respective covariance profile. c) Clinical covariance profile. 95% confidence intervals were calculated via bootstrap resampling. Note that confound removal for age, sex, education was performed prior to the PLS. d) Bootstrap ratio representing the covarying brain morphology pattern. A high positive or negative bootstrap ratio indicates high contribution of a brain region to the overall covariance profile. Vertices with a significant bootstrap ratio (> 1.96 or < -1.96) are highlighted by colors. Abbreviations: BMI – Body mass index, HDL – high density lipoprotein, LDL – low density lipoprotein, $r_{sp}$ – Spearman correlation coefficient; p – p-value; TMT-A – Trail Making Test A; TMT-B – Trail Making Test B.

#### Figure S17 – Partial least squares analysis - Hamburg City Health Study (including cognitive test results)

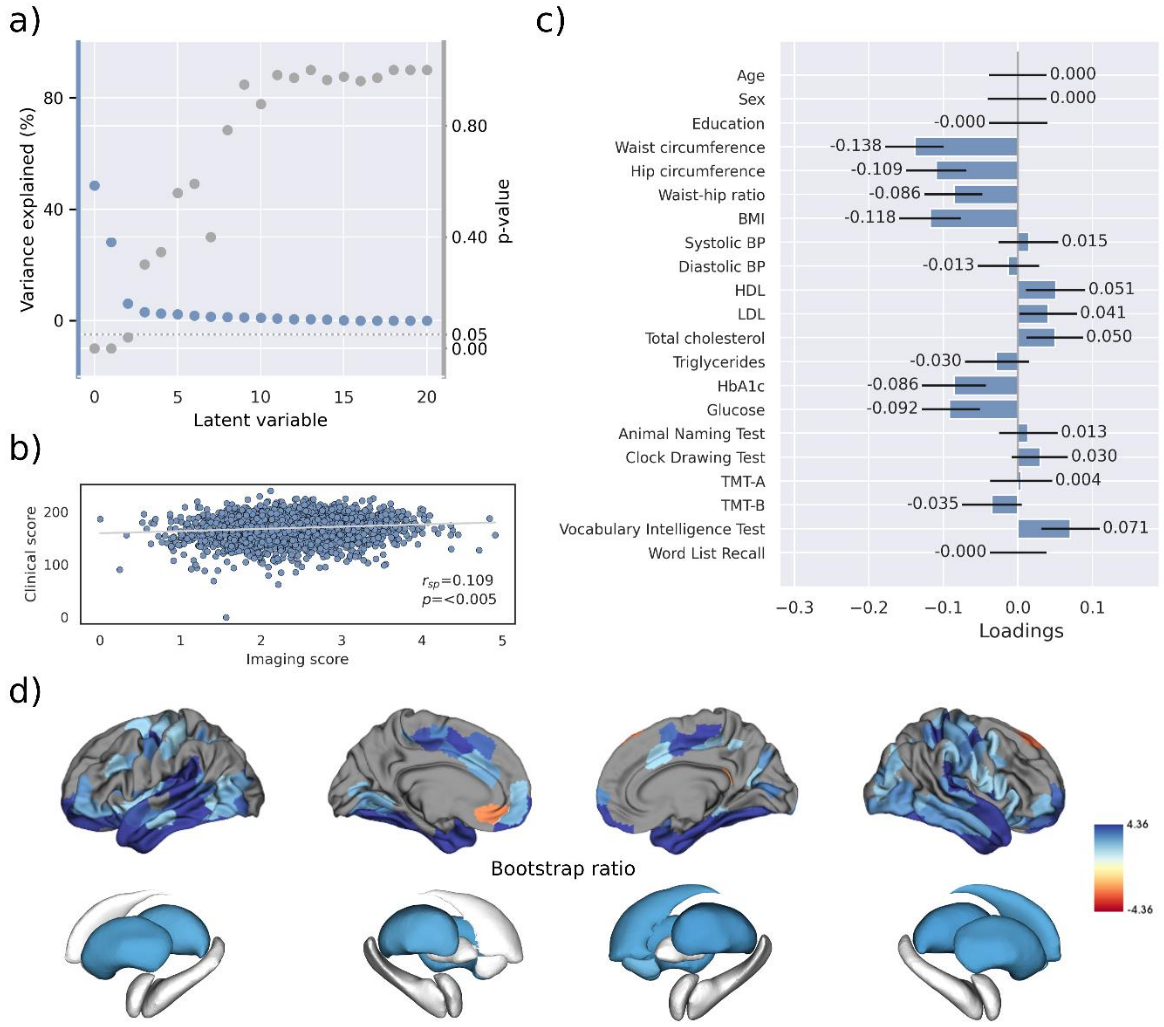

Partial least squares analysis of the HCHS subsample including cognitive test results. a) Explained variance and p-values of latent variables. b) Scatter plot relating subject-specific clinical and imaging scores. Higher scores indicate higher adherence to the respective covariance profile. c) Clinical covariance profile. 95% confidence intervals were calculated via bootstrap resampling. Note that confound removal for age, sex, education was performed prior to the PLS. d) Bootstrap ratio representing the covarying brain morphology pattern. A high positive or negative bootstrap ratio indicates high contribution of a brain region to the overall covariance profile. Vertices with a significant bootstrap ratio (> 1.96 or < -1.96) are highlighted by colors. Abbreviations: BMI – Body mass index, HDL – high density lipoprotein, LDL – low density lipoprotein, $r_{sp}$ – Spearman correlation coefficient; p – p-value; TMT-A – Trail Making Test A; TMT-B – Trail Making Test B.

### Figure S18 – Spatial correlation of effect size maps

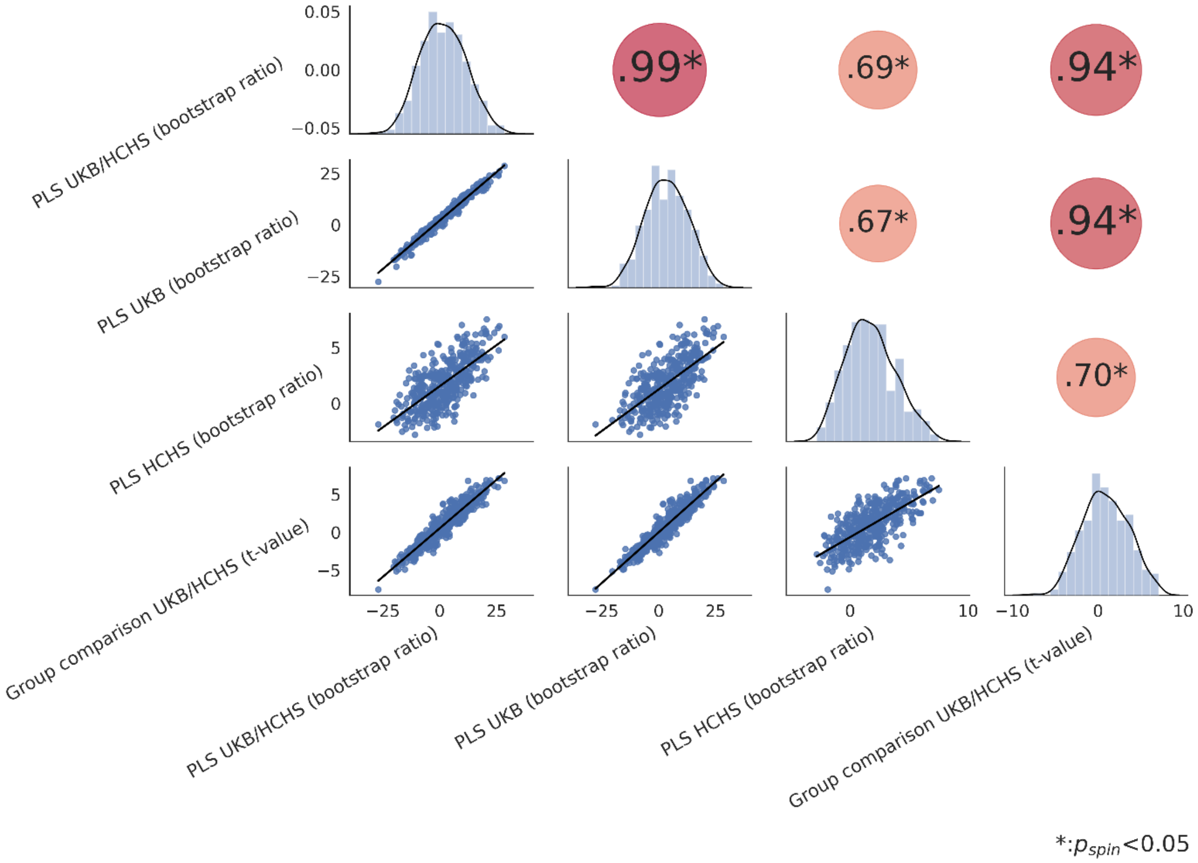

Spatial correlation matrix of all Schaefer400-parcellated metabolic syndrome effect maps (bootstrap ratio and t-statistic). The upper triangle of the matrix displays spearman correlations with dot size and color representing the magnitude of the coefficients. Asterisks highlight significant correlations after spin permutation testing and false discovery rate correction. The diagonal shows kernel density plots. The lower triangle illustrates the variables' linear relationships via regression plots. Abbreviations: HCHS – Hamburg City Health Study, PLS – Partial least squares correlation analysis; $r_{sp}$ - Spearman correlation coefficient; $p_{spin}$ – false discovery rate-corrected p-value derived from spin permutations; UKB – UK Biobank.

### Table S19 – Partial least squares analysis - Cross-validation

| **CV fold** | $\mathbf{r_{sp}}$ |
|---|---|
| 0 | 0.17 |
| 1 | 0.21 |
| 2 | 0.22 |
| 3 | 0.16 |
| 4 | 0.15 |
| 5 | 0.18 |
| 6 | 0.23 |
| 7 | 0.13 |
| 8 | 0.20 |
| 9 | 0.22 |

#### Table S20 – Virtual histology analysis – Bootstrap ratio (PLS)

| **Cell type** | **$Zr_{sp}$** | **$p_{FDR}$** |
|---|---|---|
| Endo | 0.190 | 0.016 |
| Micro | 0.271 | 0.016 |
| Ex8 | 0.165 | 0.016 |
| In1 | 0.363 | 0.036 |
| Ex6 | 0.146 | 0.034 |
| Oligo | 0.207 | 0.057 |
| In7 | 0.079 | 0.083 |
| Ex1 | 0.122 | 0.144 |
| In2 | 0.058 | 0.179 |
| In3 | 0.047 | 0.208 |
| Astro | 0.071 | 0.259 |
| In8 | 0.055 | 0.299 |
| Ex7 | 0.044 | 0.336 |
| In5 | 0.037 | 0.388 |
| Ex4 | -0.020 | 0.776 |
| Ex5 | -0.055 | 0.924 |
| In4 | -0.056 | 0.949 |
| In6 | -0.099 | 0.949 |
| Ex2 | -0.102 | 0.967 |
| Ex3 | -0.289 | 0.999 |

#### Figure S21 – Virtual histology analysis – Bootstrap ratios of latent variables 2 and 3 (PLS)

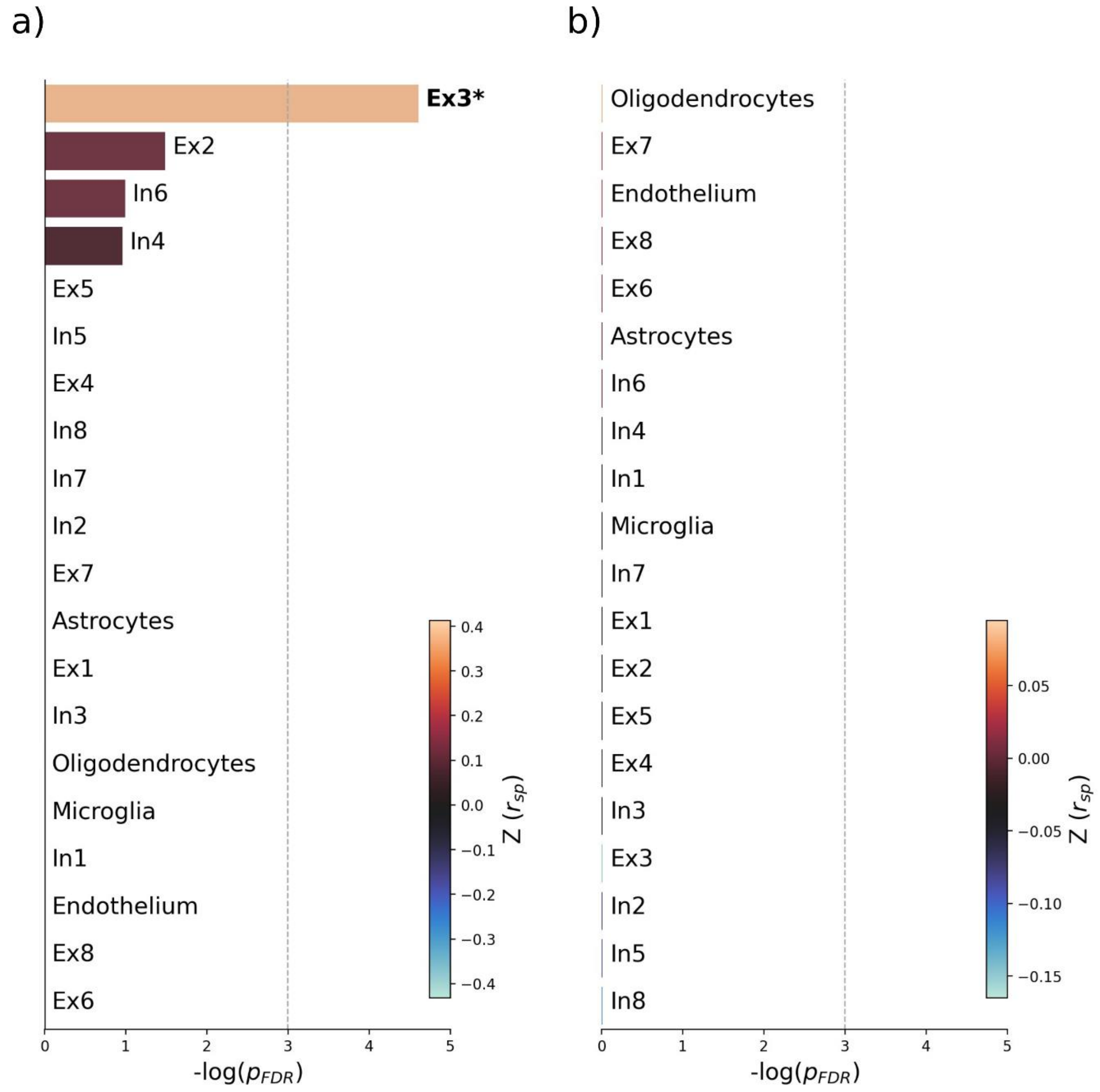

Virtual histology analysis of the bootstrap ratio maps of latent variables 2 and 3 from the PLS main analysis. Barplots display spatial correlation results of the bootstrap ratio of latent variable 2 and 3 and respective cell population densities computed via ensemble-based gene category enrichment analysis. a) Results corresponding with the bootstrap ratio of latent variable 2. b) Results corresponding with the bootstrap ratio of latent variable 3. Abbreviations: $-log(p_{FDR})$ – negative logarithm of the false discovery rate-corrected p-value derived from spatial lag models[13,14]; $r$ – Spearman correlation coeffient. $Z(r_{sp})$ – aggregate z-transformed Spearman correlation coefficient.

### Figure S22 – Virtual histology analysis – t-statistic (group comparison)

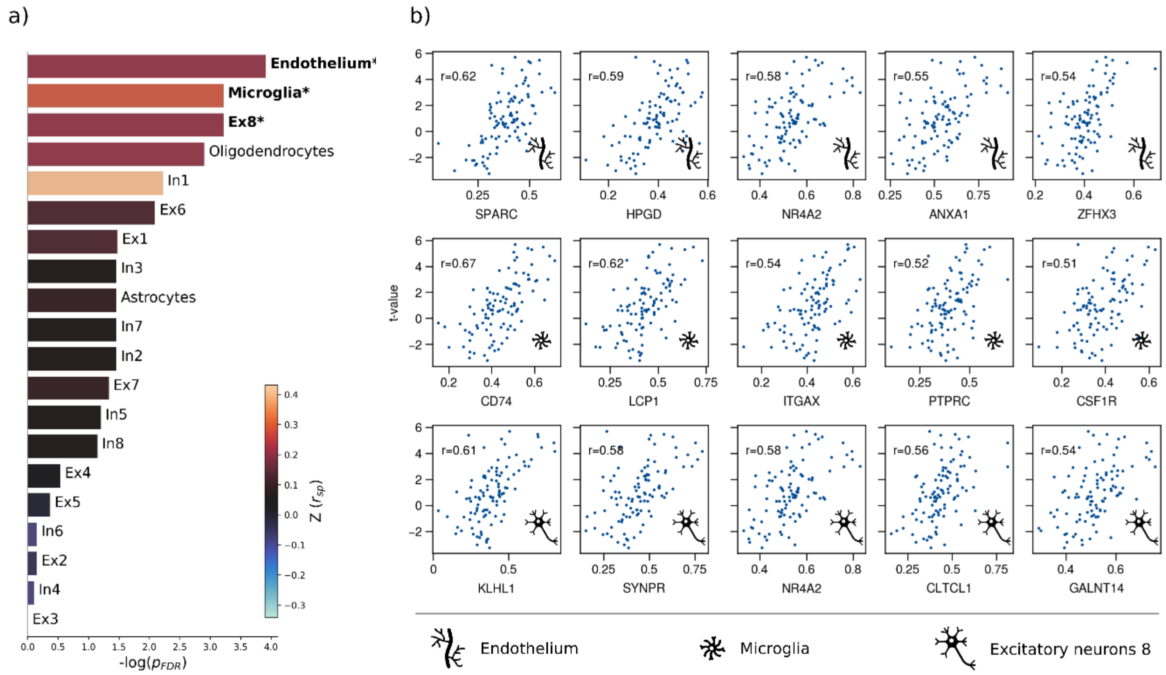

Virtual histology analysis of the t-statistic map derived from group statistics. a) Barplot displaying spatial correlation results of the bootstrap ratio and respective cell population densities computed via ensemble-based gene category enrichment analysis. b) Scatter plots illustrating per significantly associated cell population exemplary genes with top 5-highest correlation coefficients with the t-statistic map per significantly associated cell population across analyses (i.e, endothelium, microglia, excitatory neurons 8). Icons in the bottom right of each scatter plot indicate the corresponding cell type. First row: endothelium; second row: microglia; third row: excitatory neurons type 8. Abbreviations: $-log(p_{FDR})$ – negative logarithm of the false discovery rate-corrected p-value derived from spatial lag models[13,14]; $r$ – Spearman correlation coeffient. $Z(r_{sp})$ – aggregate z-transformed Spearman correlation coefficient.

### Table S23 – Virtual histology analysis – t-statistic (group comparison)

| Cell type | $Zr_{sp}$ | $p_{FDR}$ |
|---|---|---|
| Endo | 0.208 | 0.020 |
| Micro | 0.321 | 0.040 |
| Ex8 | 0.208 | 0.040 |
| Oligo | 0.233 | 0.055 |
| In1 | 0.432 | 0.108 |
| Ex6 | 0.145 | 0.123 |
| Ex1 | 0.156 | 0.229 |
| In3 | 0.058 | 0.233 |
| Astro | 0.120 | 0.233 |
| In7 | 0.059 | 0.233 |
| In2 | 0.063 | 0.233 |
| Ex7 | 0.089 | 0.263 |
| In5 | 0.063 | 0.300 |
| In8 | 0.066 | 0.317 |
| Ex4 | 0.015 | 0.585 |
| Ex5 | -0.007 | 0.690 |
| In6 | -0.078 | 0.861 |
| Ex2 | -0.070 | 0.861 |
| In4 | -0.087 | 0.901 |
| Ex3 | -0.341 | 0.997 |

### Figure S24 – Network contextualization – t-statistic (group comparison)

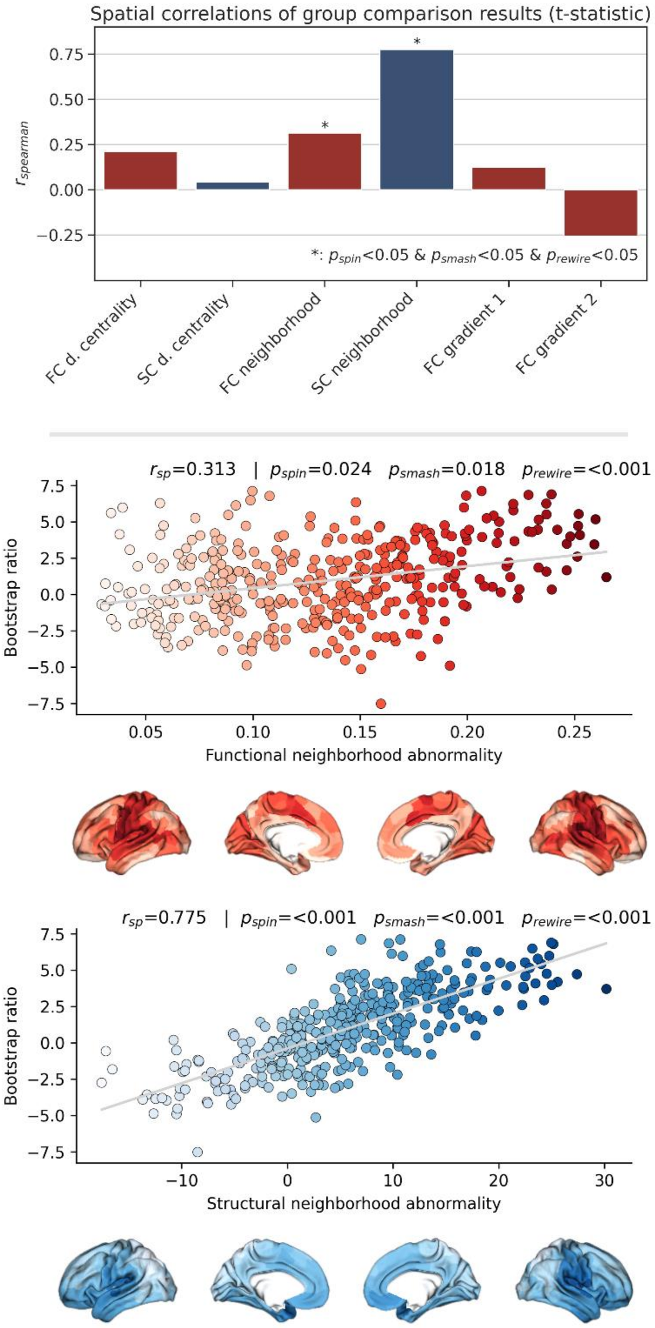

Brain network contextualization analysis of group statistics results. Results are presented for t-statistics maps derived from group statistics considering the pooled sample of UK Biobank subjects and HCHS subjects. The upper row barplot summarizes the analysis results displaying the Spearman correlation with regard to each investigated index. Asterisks indicate statistical significance with respect to spin, brainSMASH and network rewiring null models. The middle and lower row display scatter plots of the significant association of the t-statistics map and the functional and structural neighborhood abnormality, respectively. The scatter plots are supplemented by surface plots for anatomical localization.

### Figure S25 – Network contextualization with Hamburg City Health Study connectomes

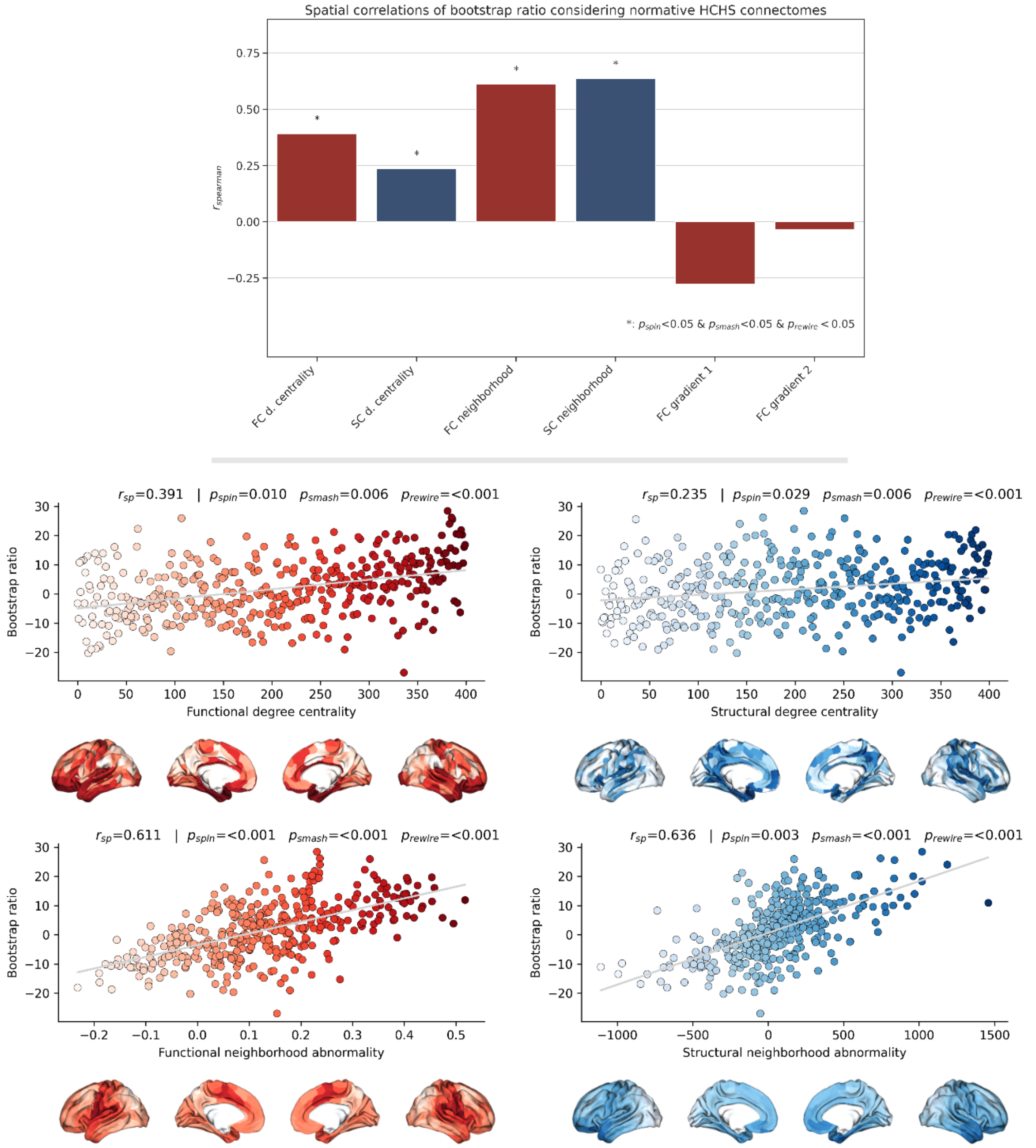

Brain network contextualization analysis of partial least squares correlation results (bootstrap ratio) based on group-consensus connectomes from the Hamburg City Health Study. Results are presented for bootstrap ratio maps derived from partial least squares correlation analysis considering the pooled sample. The upper row bar plot summarizes the analysis results displaying the Spearman correlation with regard to each investigated index. Asterisks indicate statistical significance with respect to spin, brainSMASH and network rewiring null models. Scatter plots that illustrate the significant spatial relationships are presented below. The middle row displays the relationship of the bootstrap ratio map and the ranked functional and structural degree centrality. The lower row illustrates the association of the bootstrap ratio map and the functional and structural neighborhood abnormality.

### References

1. Alberti, K. G. M. M., Zimmet, P. & Shaw, J. Metabolic syndrome--a new world-wide definition. A Consensus Statement from the International Diabetes Federation. *Diabet Med* **23**, 469–480 (2006).
2. Larivière, S. *et al.* The ENIGMA Toolbox: multiscale neural contextualization of multisite neuroimaging datasets. *Nat Methods* **18**, 698–700 (2021).
3. Larivière, S. *et al.* Network-based atrophy modeling in the common epilepsies: A worldwide ENIGMA study. *Sci. Adv.* **6**, eabc6457 (2020).
4. van den Heuvel, M. P. & Sporns, O. Network hubs in the human brain. *Trends in Cognitive Sciences* **17**, 683–696 (2013).
5. Rubinov, M. & Sporns, O. Complex network measures of brain connectivity: uses and interpretations. *Neuroimage* **52**, 1059–1069 (2010).
6. Shafiei, G. *et al.* Spatial Patterning of Tissue Volume Loss in Schizophrenia Reflects Brain Network Architecture. *Biological Psychiatry* **87**, 727–735 (2020).
7. Petersen, M. *et al.* Brain network architecture constrains age-related cortical thinning. *NeuroImage* **264**, 119721 (2022).
8. Margulies, D. S. *et al.* Situating the default-mode network along a principal gradient of macroscale cortical organization. *Proc Natl Acad Sci U S A* **113**, 12574–12579 (2016).
9. Sydnor, V. J. *et al.* Neurodevelopment of the association cortices: Patterns, mechanisms, and implications for psychopathology. *Neuron* **109**, 2820–2846 (2021).
10. Vos de Wael, R. *et al.* BrainSpace: a toolbox for the analysis of macroscale gradients in neuroimaging and connectomics datasets. *Communications Biology* **3**, 1–10 (2020).
11. Ho, D., Imai, K., King, G. & Stuart, E. A. MatchIt: Nonparametric Preprocessing for Parametric Causal Inference. *Journal of Statistical Software* **42**, 1–28 (2011).

12. Larivière, S. *et al.* BrainStat: A toolbox for brain-wide statistics and multimodal feature associations. *Neuroimage* **266**, 119807 (2023).

13. Dukart, J. *et al.* JuSpace: A tool for spatial correlation analyses of magnetic resonance imaging data with nuclear imaging derived neurotransmitter maps. *Hum Brain Mapp* **42**, 555–566 (2021).

14. Burt, J. B. *et al.* Hierarchy of transcriptomic specialization across human cortex captured by structural neuroimaging topography. *Nat Neurosci* **21**, 1251–1259 (2018).
